# *Aspergillus nidulans* Transcription Factor BrlA is Utilized in a Conidiation-Independent Response to Cell-Wall Stress

**DOI:** 10.1101/2024.11.21.624663

**Authors:** Harley Edwards, Joseph Zavorskas, Walker Huso, Alexander G. Doan, Kelsey Grey, JungHun Lee, Meredith Morse, Heather H. Wilkinson, Danniel Ebbole, Brian D. Shaw, Steven D. Harris, Ranjan Srivastava, Mark R. Marten

## Abstract

Under synchronized conidiation, over 2500 gene products show differential expression, including transcripts for both *brlA* and *abaA*, which increase steadily over time. In contrast, during wall-stress induced by the echinocandin micafungin, the *brlA* transcript is upregulated while the *abaA* transcript is not. In addition, when *mpkA* (last protein kinase in the cell wall integrity signaling pathway) is deleted, *brlA* expression is not upregulated in response to wall stress. Together, these data imply BrlA may play a role in a cellular stress-response which is independent of the canonical BrlA-mediated conidiation pathway. To test this hypothesis, we performed a genome-wide search and found 332 genes with a putative BrlA response element (BRE) in their promoter region. From this set, we identified 28 genes which were differentially expressed in response to wall-stress, but not during synchronized conidiation. This set included seven gene products whose homologues are involved in transmembrane transport and 14 likely to be involved in secondary metabolite biosynthesis. We selected six of these genes for further examination and find that they all show altered expression behavior in the *brlA* deletion strain. Together, these data support the idea that BrlA plays a role in various biological processes outside asexual development.

**Importance:** The *Aspergillus nidulans* transcription factor BrlA is widely accepted as a master regulator of conidiation. Here, we show that in addition to this function BrlA appears to play a role in responding to cell-wall stress. We note that this has not been observed outside *A. nidulans*. Further, BrlA-mediated conidiation is highly conserved across *Aspergillus* species, so this new functionality is likely relevant in other *Aspergilli*. We identified several transmembrane transporters that have altered transcriptional responses to cell-wall stress in a *brlA* deletion mutant. Based on our observation, together with what is known about the *brlA* gene locus’ regulation, we identify *brlAβ* as the likely intermediary in function of *brlA* in the response to cell-wall stress.

## Introduction

The filamentous fungus *Aspergillus nidulans* is a model organism that has tremendous relevance in biomanufacturing (*1*), bioremediation (*2*), and medicine (*3, 4*). *A. nidulans* shows significant similarity to *A. oryzae* and *A. niger* which have long been used in industrial applications (e.g., fermenting soy sauce and citric acid (*1*)). *A. nidulans* also serves as a model for *Aspergillus spp*. deemed “hyperaccumulators,” of interest for their ability to metabolize or sequester heavy metals, petroleum hydrocarbons, synthetic dyes, and agricultural chemicals (*2*). *A. nidulans* also has significant similarity to pathogenic fungi such as *A. fumigatus* (*3*), and thus provides an effective system for pathogenic mechanisms of action (*4*). Thus, an increased understanding of *A. nidulans* morphology, physiology, and developmental processes has potential to control fungal growth and disease. Of particular importance is developing a robust understanding of asexual reproduction in *Aspergillus spp*.

Conserved across *Aspergillus spp*. is a central, asexual reproduction pathway of three transcription factors: BrlA, AbaA, and WetA (*5, 6*). All three play essential, temporal and spatial roles in successful asexual reproduction (*6, 7*). For example, the pathway is known to activate the genes in order: *brlA, abaA, wetA* (*8, 9*). Also, it has been shown that (i) loss of *brlA* leads to a complete loss of conidiophore development (*8*), (ii) loss of *abaA* confers a loss of phialide differentiation (*10*), and (iii) loss of *wetA* leads to loss of pigment, reduced hydrophobicity, and reduced viability of fully formed spores (*11*).

While this gene expression cascade has been widely studied in the context of developmental morphology, contemporary work has discovered additional physiological roles that vary across *Aspergillus spp.* For example, in *A. fumigatus brlA* has been shown to affect gliotoxin biosynthesis(*12*), and other tissue-specific secondary metabolites(*13*). Expression of *brlA* is also affected by secondary metabolism in *A. oryzae(14)*, and *A. flavus(15).* In *A. nidulans*, *brlA* plays a role during both autophagy (*16*) and autolysis (*17*). In addition, other regulators that control expression of *brlA* (e.g., *laeA*(*18*) and *rocA*(*19*)) also control secondary metabolite production. More recently, it has been noted that *A. nidulans* increases expression of *brlA* in response to echinocandin-mediated, cell-wall stress (*20, 21*). However, the scope and nature of the function of BrlA outside of its canonical role in asexual development remains poorly understood.

In *A. nidulans*, the *brlA* locus has two overlapping transcription units: *brlAα* and *brlAβ* (*22*). The nucleotides coding for *brlAβ* overlap *brlAα*, and code for 23 more amino acids at the N-terminus of the resulting peptide (*23*). BrlAα and BrlAβ are individually essential for proper conidial development, as a mutant lacking either subunit causes distinct morphological differences in how conidiation is interrupted (*22*). Despite being independently essential, the over-expression of either subunit can lead to the restoration of normal development (*22, 23*). The fact that overexpression of either subunit can restore function implies the different subunits are not distinct in protein activity, but are distinct in how their spatiotemporal distribution coordinates various roles at different stages of growth (*22*). The *brlA* locus also contains a micro-ORF (µORF), upstream of the coding region of the *brlAβ* subunit (*23*). It has been shown that the µORF inhibits premature development of conidiation in submerged culture by inhibiting translation of BrlAβ (*23*).

In previous work, we used RNA-seq to characterize the dynamic transcriptome of *A. nidulans* in response to micafungin-induced cell-wall stress (*20, 21*). Under micafungin exposure, *brlA* is one of the top 50 most significantly overexpressed transcripts (*20*). Interestingly, *abaA* and *wetA* do not show the same timing, or the same magnitude of expression, as they do during normal asexual development. Additionally, we found that a key cell-wall stress mediator, MpkA, likely interacts with BrlA in response to wall stress (*21*). This Cell Wall Integrity (CWI) to BrlA connection has also been hypothesized by others in the past (*24, 25*). Our findings here demonstrate that the significant upregulation of *brlA* during wall stress does not occur in an *mpkA* deletion (Δ*mpkA*) strain. Due to this apparent short-circuiting of the asexual reproduction pathway, we hypothesize that some expressed subunit of the *brlA* locus is involved in the MpkA mediated cell-wall stress response in a manner independent of the well-known *brlA, abaA, wetA* pathway for conidiation.

To test this hypothesis, we utilized dynamic RNA-seq data describing *A. nidulans* under synchronized conidiation conditions, coupled with dynamic transcriptomic data describing *A. nidulans* response to micafungin exposure. We used the Derivative Profiling omics Package (DPoP)(*26*) to analyze the dynamics of both of these data sets to identify genes that are differentially regulated under each condition. We then used prior knowledge of the BrlA recognition elements (BREs) (*27*) to conduct a genome-wide motif search for genes with BREs in their upstream region. By comparing these groups, we discovered evidence that BrlA regulates 28 genes in a conidiation-independent response to cell wall stress. The resulting genes are highly correlated with one another, yielding 4 distinct groups: transmembrane transporters, secondary metabolite regulation, DNA modulation, and a fourth group of unknown function. We follow up on this evidence with qPCR experiments to determine if transmembrane transporter expression is significantly affected in a *brlA* mutant. Our work provides compelling evidence for a previously unrecognized role of BrlA and its µORF in the response to cell wall stress.

## Results

Initial work involved determining how regulatory genes involved in asexual-development change expression in response to synchronized conidiation. We induced synchronous conidiation in *A. nidulans* at the air/liquid interface as described in the methods section. Samples were then removed over 48 hours, and subjected to RNA-seq to assess transcript expression (**Figure 1**). We found both *brlA* and *abaA* transcripts increased significantly over time, showing 1000-fold and 10-fold increases respectively. In contrast, *wetA* and the *brlA* μORF showed relatively little change in expression over time.

**Figure 1.**
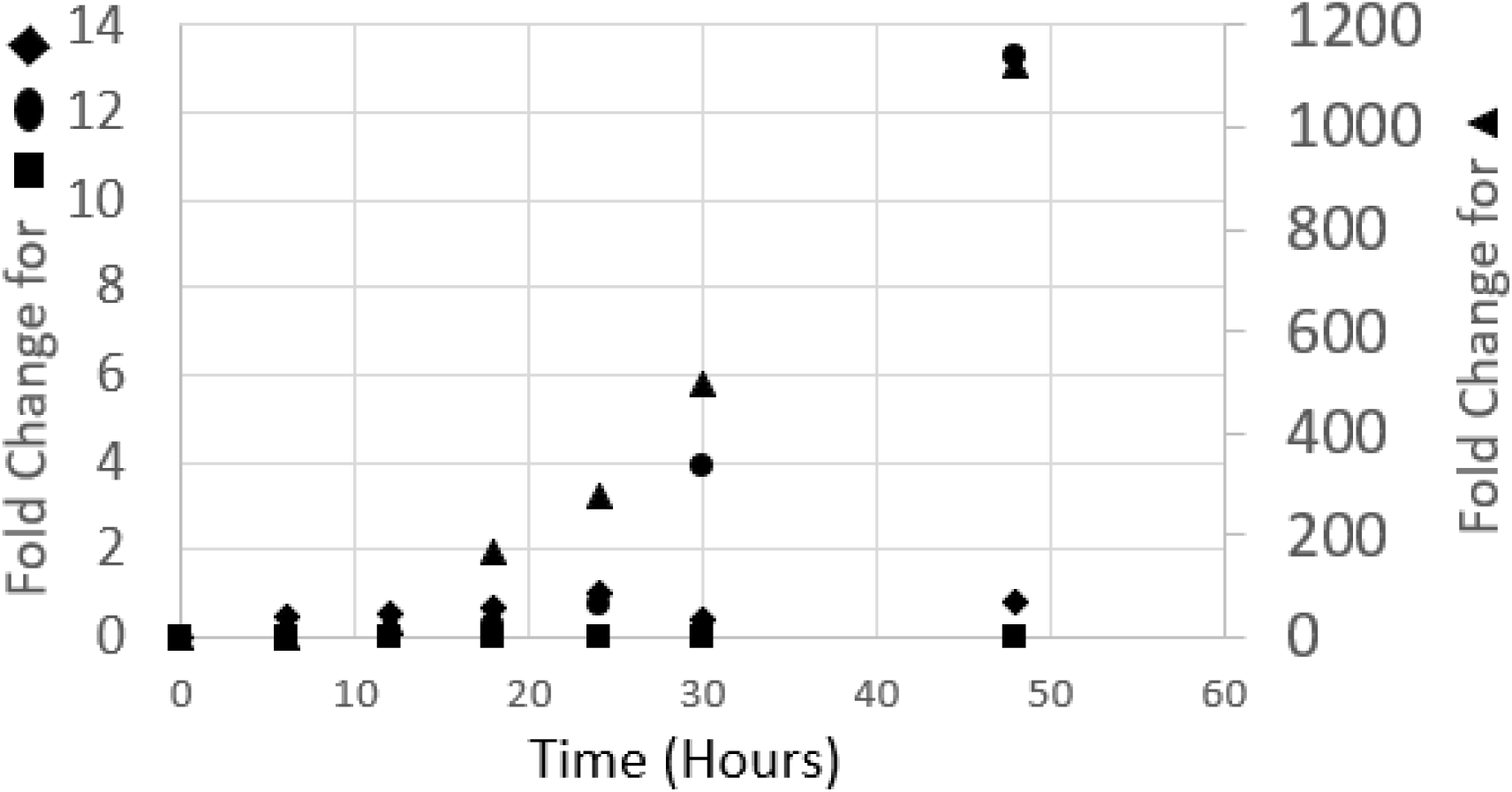
Change in transcript expression of asexual-development regulatory genes in Aspergillus nidulans during conidiation. Symbols: ▴ brlA; ● abaA; ◆ wetA; ▪ brlA μORF. Fold change is the difference between the transcript-length normalized counts from the first and reference timepoint, divided by the first timepoint.

We have also studied the *A. nidulans* response to cell-wall stress (21, 22). To do this, we grew *A. nidulans* in flasks and added the echinocandin micafungin during exponential growth. We then removed samples over time (2h) and subjected them to RNA-seq to assess transcript expression (**Figure 2**). Unexpectedly, we found changes in expression of asexual-development regulatory genes. For example, we found that both *brlA* and the *brlA* μORF transcripts are dynamically-upregulated in response to micafungin exposure, increasing approximately 10-fold over 2h exposure. In contrast, *abaA* and *wetA* transcripts show relatively little change in expression level. Further, we determined that increased expression of both *brlA* and the *brlA* μORF is mediated by a functional copy of the *mpkA* gene, as the Δ*mpkA* strain did not show the same upregulation of *brlA* and μORF transcripts during wall stress.

**Figure 2.**
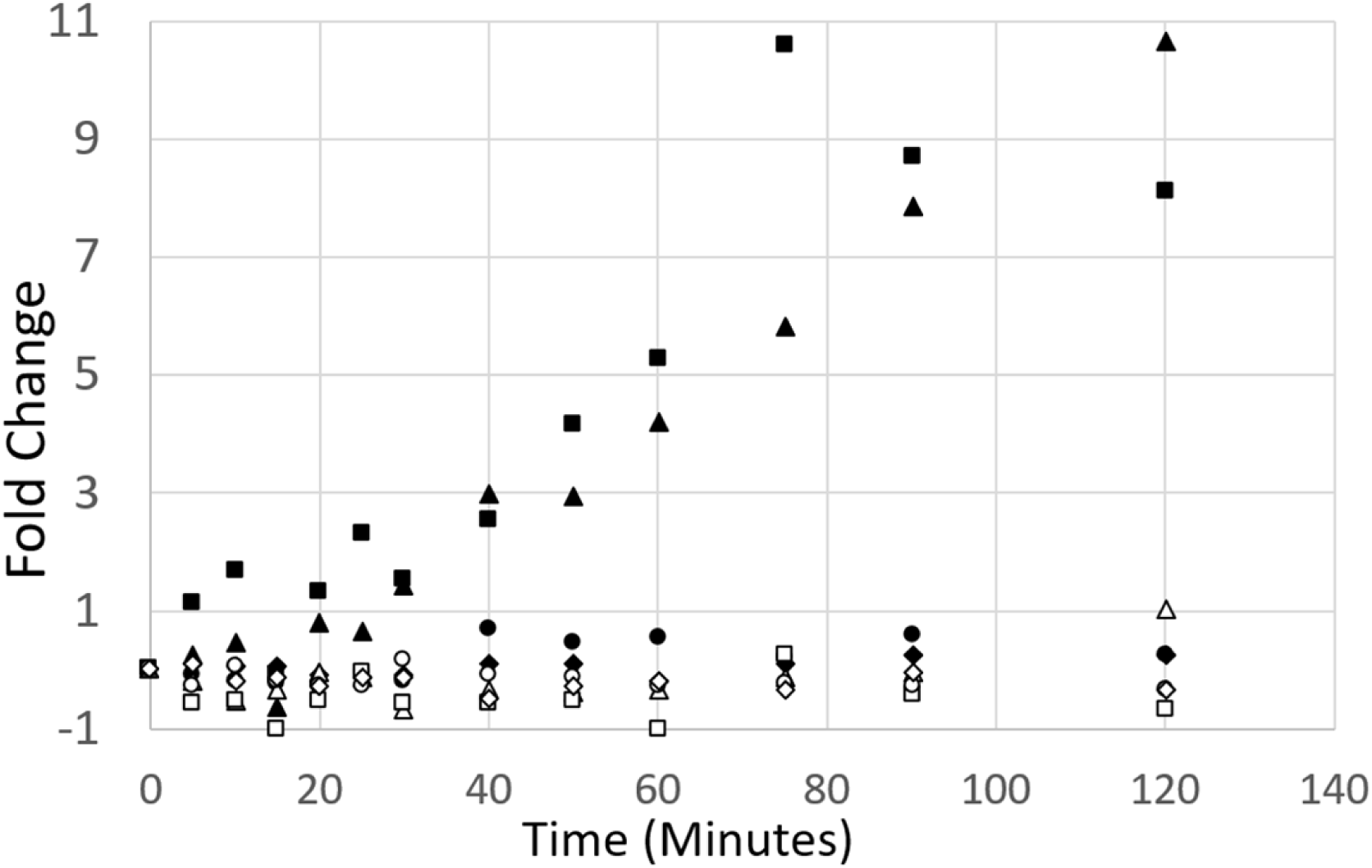
Change in transcript expression of asexual-development regulatory genes, in mpkA+/- strains, after exposure to micafungin. Micafungin added to exponentially growing A. nidulans cultures at time zero. Black shapes from mpkA+ strain (A1405;(20)), white shapes mpkA-strain (A1404;(21)). Symbols: ▲,△ brlA; ●,◯ abaA; ◆,◇ wetA; ▪,□ brlA μORF. Fold change is the difference between the transcript-length normalized counts from the first and reference timepoint, divided by the first timepoint. Data show only brlA and the μORF have increased expression in response to cell wall stress, and only in the mpkA+ strain.

Comparing the responses in Figures 1 and 2, implies that *A. nidulans* utilizes the transcription factors *brlA* and the *brlA* μORF differently during wall-stress than during asexual development. Furthermore, the wall stress response appears to be mediated by the well-studied cell wall integrity (CWI) signaling pathway, where MpkA is the final protein kinase.

To further assess these different responses, we subjected both of these transcriptomic data sets to analysis via DPoP(*26*) to identify significant, differentially-expressed transcripts under each condition (i.e., synchronized conidiation and wall stress). Results are shown in **Figure 3**. We find that during synchronized conidiation 2730 genes are differentially regulated (Figure 3, pink circle), while during cell-wall stress 1839 genes are differentially expressed (Figure 3, blue circle). Comparing these conditions, we find 927 genes showed differential expression in response to wall stress but not during asexual development. This is particularly useful considering that *brlA* is uniquely upregulated in response to micafungin whereas *abaA* and *wetA* are not.

**Figure 3.**
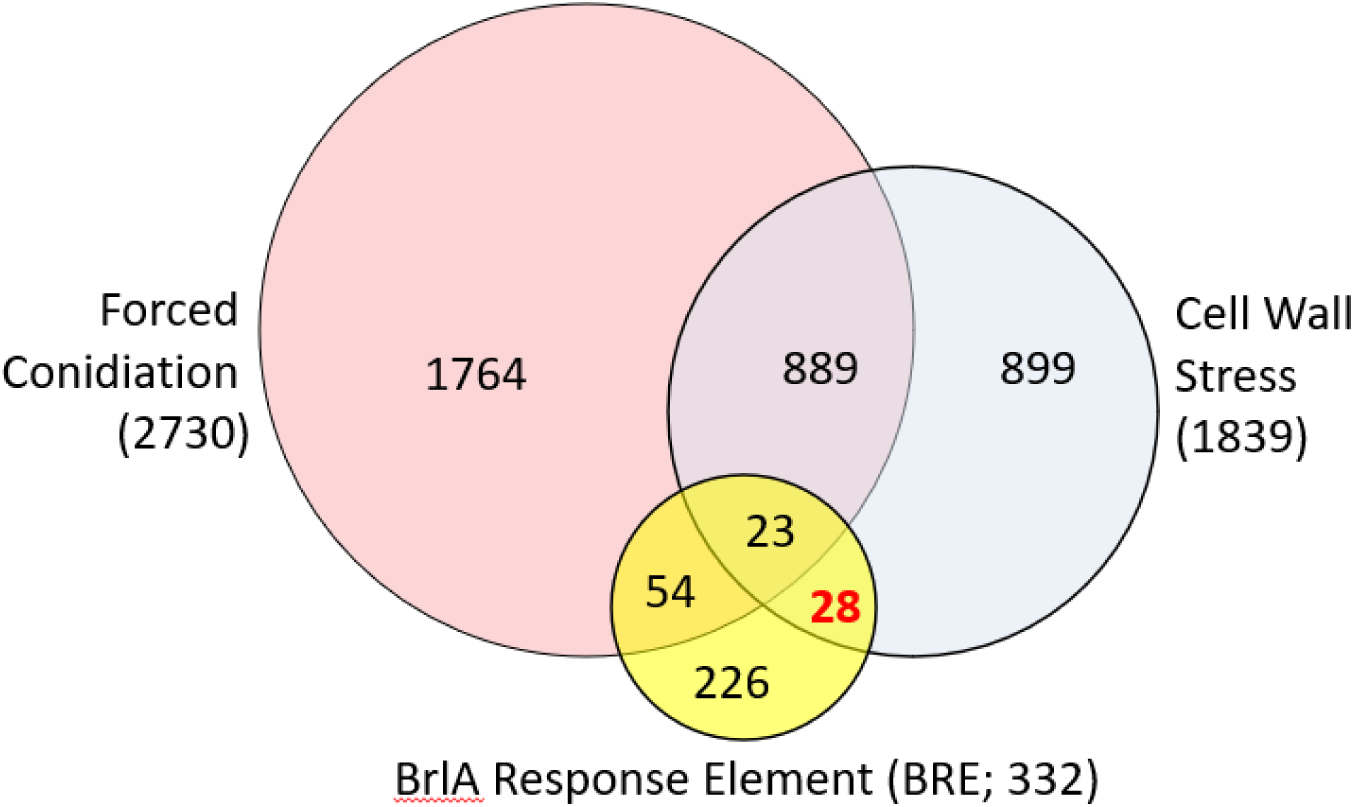
Venn Diagram Comparison of differentially expressed genes and those with BrlA response element (BRE).

To determine which of these 927 genes may have been specifically regulated by BrlA, we utilized the DNA sequence where BrlA binds, known as the BrlA response element (BRE) (27). This bioinformatic search is described in the methods section and code to reproduce the analysis is in supplementary file (S1). We searched the *A. nidulans* genome for genes downstream of the BRE, and identified 332 (Figure 3, yellow circle) candidates (Supplementary File S2). When compared to the list of 927 genes, we find 28 genes downstream of a BRE, expressed in a conidiation-independent response to cell wall stress. These 28 genes and their predicted functions are shown in Table 1.

**Table 1:** Conidiation-independent micafungin response genes with upstream BREs.

| Gene | Name | Homologue function | Trend (micafungin) |
| --- | --- | --- | --- |
| Transmembrane Transporters |  |  |  |
| AN8595 |  | MFS efflux pump (multidrug resistance) | Up |
| AN9165 |  | MFS efflux pump (multidrug resistance) | Up, Down |
| AN3247 |  | ABC transporter (multidrug resistance) | Up |
| AN8610 |  | MFS efflux pump (multidrug resistance) | Up |
| AN3304 |  | GABA transporter (drug resistance) | Up |
| AN0010 |  | Neutral amino acid transporter | Down |
| Secondary Metabolite Biosynthesis |  |  |  |
| AN0009 |  | 5-oxoprolinase | Down |
| AN5311 |  | Tyrosinase/catechol oxidase | Down, Up |
| AN5616 |  | kynurenine aminotransferase | Up |
| AN2834 |  | Putative rhamnogalacturonan acetyl esterase | Down, Up |
| AN7860 |  | beta lactamase domain containing protein | Down |
| AN7541 | **cut2** | Cutinase/lipidase/esterase | Down |
| AN7828 | **urhB** | Rhamnogalacturonan hydrolase | Up, Down |
| AN7836 |  | small cysteine-rich protein | Down, Up |
| AN8330 |  | PKS_ER domain containing | Down, up |
| AN8155 |  | NADPH-dependent "short chain dehydrogenase" | Up |
| AN7075 |  | FAD/NAD binding; redox; environmental adaptation | Up, Down |
| AN6847 |  | aryl sulfatase/alkaline phosphatase | Up |
| AN5977 |  | ketoreductase; NADP/NADPH as acceptor | Up |
| AN4975 |  | FAD/NAD binding redox protein | Up, Down |
| DNA Binding |  |  |  |
| AN5929 |  | con7, master regulator of virulence morphology, essential for conidiation, zinc finger, CH2H domain containing | Down |
| AN7580 |  | unknown CH2H transcription factor | Up |
| AN3243 |  | Methyltransferase, "yukJ" ortholog | Up, Down |
| AN0621 |  | long interspersed nuclear element (LINE) [reverse transcriptase] | Up, Down |
| Unknown |  |  |  |
| AN1378 |  | calcium regulation protein, SMP-30 | Down, Up |
| AN5385 |  | Unknown | Up |
| AN3553 |  | Unknown | Up, Down |
| AN6011 |  | Unknown | Down |

To validate these observations, we assessed transcription of *brlA*, *abaA*, *wetA,* and the six transmembrane transporters shown in Table 1. To conduct these tests, we used qPCR to carry out transcriptional analysis of *brlA* +/- strains, both with and without micafungin (MF +/-). For each of the genes of interest, a significant change in the response to micafungin under loss of *brlA* can be observed in Figure 4.

**Figure 4.**
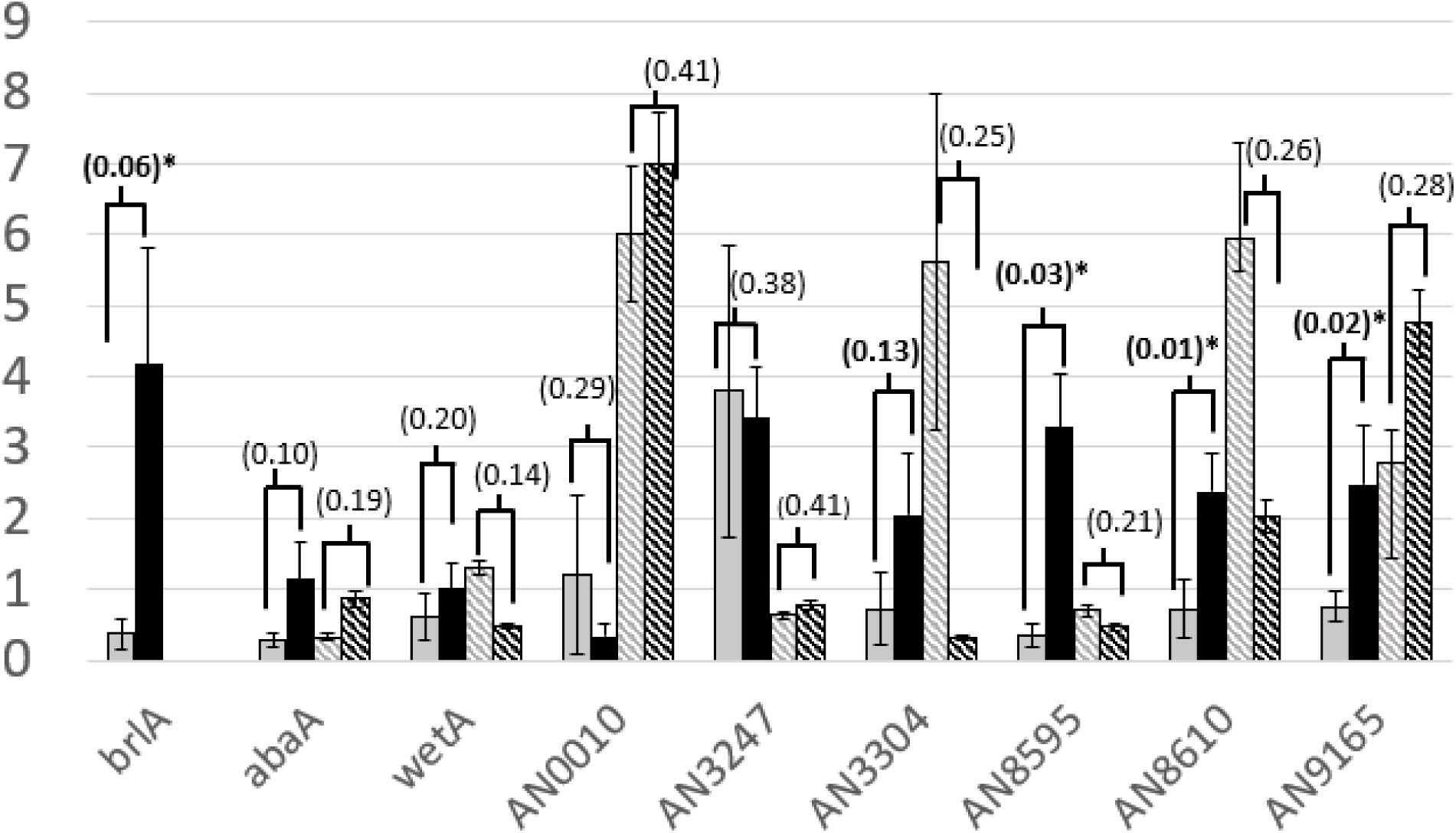
Altered gene expression one hour after micafungin exposure. For each gene listed, bars represent data from brlA+ strain (control; A1405) with no micafungin (solid gray), and 10ng/mL micafungin treatment (solid black). These are followed by data from a brlA-deletion strain (A1826), with no micafungin (hatched gray) and 10ng/mL micafungin (hatched black). Bars represent mean, relative, fold-change of the efficiency-corrected starting concentration, (ΔΔN_0_), for the target gene relative to histone after one hour. All data collected from RNA extractions in mid-exponential phase growth, at the time of perturbation, and again one hour later. Two statistical significance values are shown for each set of bars. First p-value compares data for brlA+ strain (A1405) without and with micafungin. Second p-value compares data for brlA-deletion strain (A1826) with and without micafungin. Error bars are standard error with n=4 for all data.

The difference in the control (i.e., *brlA*+) strain response to micafungin for the transcripts AN8595, AN8610, and AN9165 was statistically significant at the 95% confidence interval. All six transcript trends agree with the trend observed previously in the RNAseq data, (Table 1; Trend). In the *brlA* deletion mutant (i.e., Δ*brlA*) the overall trend is altered for every transcript. Any transcript which showed a statistically significant difference in the control strain, does not undergo a statistically significant difference in expression in the Δ*brlA* strain.

## Discussion

Our work implies that the *A. nidulans brlA* transcription factor does not act only as the master regulator of conidiation but may play a role in the echinocandin cell-wall stress response. We show that: (i) MpkA mediates a signal to upregulate *brlA* in response to cell wall stress. (ii) There is a difference in expression dynamics for this regulatory system under the two differing conditions of cell wall stress or conidiation. (iii) Analysis of multiple transcriptomic data sets reveals genes of interest that are likely controlled in some capacity by BrlA (i.e., genes which were differential in response to micafungin, were not differential during conidiation, and that had BREs in their promoter region. (iv) Several genes of interest have significantly altered transcriptional activity in response to cell-wall stress in a *brlA* deletion mutant, including three transmembrane transporters (AN8595, AN8610 and AN9165) which are potentially involved in a protective response to micafungin.

We note the difference in expression dynamics for the *brlA* μORF*, brlA, and abaA* under synchronized conidiation, where the *brlA* μORF is not expressed but *brlA* and *abaA* are. In contrast, under micafungin treatment, expression of the *brlA* μORF is nearly identical to that of *brlA* and *abaA* is not expressed. This difference is consistent with known behavior of AbaA as a co-regulator of conidiation. This finding is also consistent with prior work showing the *brlA* μORF gene product is a repressor of asexual development (*23*). Taken together, our work implies that the conidiation-independent cell-wall stress response of the *brlA* locus control system utilizes the *brlA* μORF to inhibit further progression of the conidial pathway which would express/utilize *abaA*. Our findings suggest that the *brlA* μORF stops normal development under cell wall stress and, based on prior work, initiates micro-cycle conidiation (*21*). Since *brlA* μORF inhibits *brlAβ* and not *brlAα*, this suggests that *brlAα* plays some role in the *brlA* mediated response to cell-wall stress.

### The brlA μORF in the echinocandin mediated cell wall stress response

It has been shown that expression of the *brlA* μORF represses both asexual development and *abaA* expression through translationally inhibiting *brlAβ*, but not *brlAα* (*23*). We show that the *brlA* μORF is not expressed during normal asexual development (Figure 1) but is expressed during cell-wall stress (Figure 2). These expression dynamics suggest that the *brlA* μORF repression of asexual development is occurring during the cell-wall stress response. We hypothesize that this acts as a checkpoint that allows *brlA*α to serve its function but stops *brlAβ* activation of *abaA*. This includes BrlAα activation of genes listed in this work (transmembrane transporters), but also BrlAα activation of genes excluded from our list of genes (differential during conidiation), which may play a role in micro-cycle conidiation observed to occur under cell wall stress at these conditions(*21*).

### MpkA mediates the cell wall stress response of the brlA μORF

MpkA has been shown to physically associate with BrlA and to affect asexual development (*24*). The terminal protein in the CWI cascade, transcription factor RlmA, has been shown to transduce a signal to BrlA(*25*). Previous work has also shown that micro-cycle conidiation in response to cell wall stress does not occur in an *mpkA* mutant (*21*). Here we show that the stress response of the *brlA* locus, specifically the increased expression of *brlA* or the *brlA* μORF, is not observed in an *mpkA* mutant. We interpret these findings as compelling evidence that the CWI signaling pathway mediates the response to cell wall stress responsible for *brlA* μORF expression.

### brlA, abaA and wetA have drastically different dynamic ranges

These three major regulatory transcription factors, *brlA*, *abaA* and *wetA* have drastically different dynamic ranges in expression. It has been shown that *brlA* is one of the 50 most upregulated genes under micafungin exposure(*20*), and is also in the top 50 most upregulated genes by fold change in the dynamic, transcriptomic, synchronized conidiation data set presented here. Over the 48 hours of the synchronized conidiation experiment, *brlA* was upregulated over one-thousand-fold, while over the 2 hours of micafungin treatment it is upregulated 10-fold. This implies *brlA* has a very large dynamic range and is utilized at various expression levels for its multiple distinct functions.

Additionally, *abaA* is only 10-fold upregulated over the 48 hours of conidiation, and never greater than 1-fold change under micafungin treatment. This evidence suggests that something outside of the known *brlA, abaA, wetA* pathway must either repress *abaA* under cell wall stress or activate *abaA* under conidiation. Our data suggests repression of *abaA* activation during cell-wall stress. This is due to *brlA* μORF expression seen at the same time which would inhibit translation of *brlAβ* and subsequently *abaA* expression. We also note the lack of any dynamics in the expression of *wetA*. Despite the difference in time or the difference in conditions subjected to the cells, *wetA* is never upregulated in a comparable manner to *brlA* and *abaA*. This demonstrates that *wetA* has the lowest dynamic range of any of genes in the pathway.

### Downstream gene control by brlA

qPCR was used to examine the relative gene expression of six genes of interest after micafungin perturbation in both a control strain (A1405) and the Δ*brlA* mutant (A1826). Of the genes tested via qPCR, all displayed altered gene expression in the mutant strain. These six were AN0010, AN3247, AN3304, AN8595, AN8610, and AN9165. The latter three, AN8595, AN8610 and AN9165, show a statistically significant increase in gene expression in response to micafungin treatment in the control strain, whereas in the *brlA* mutant the statistically significant upregulation in response to micafungin is not observed. In AN0010, AN3247 and AN3304, the gene expression in response to micafungin in the control strain agrees with the trend predicted to occur, but expression in the *brlA* deletion mutant changes overall trend and magnitude. All these genes are homologues of transmembrane efflux pumps with roles in antibiotic/antitoxin resistance with the exception of AN0010 which is the homologue of an amino acid permease. We hypothesize that *brlAα* is responsible for the increased gene expression of these transmembrane efflux pumps, since *brlAβ* is likely translationally inhibited by the *brlA* µORF during cell wall stress.

### Other Effects of Wall Stress

In the list of 28 genes of interest (Table 1), other than transmembrane transporters, there were genes related to secondary metabolites, and the redox potential molecules to support that kind of advanced polyketide synthesis (Table 1). This is consistent with recent findings that *brlA* regulates secondary metabolite biosynthesis in various *Aspergillus* species (*12-15*). Also, the third group of well described genes in the 28 are annotated as DNA modulation and regulation genes, exactly the description expected from a transcription factor (BrlA) which controls other transcription factors. Between our qPCR confirmatory experiments, recent findings about the role of BrlA in secondary metabolism in other *Aspergilli spp.*, and the orthology/homology description of the set of genes matching transcriptional activity, we have high confidence in the relevance of the genes found in our analysis.

### BRE Related Genes

Our analysis reveals other interesting patterns of gene expression relative to conidiation and cell wall stress responses. For example, of the 332 genes which have a BRE, almost two-thirds, 226 genes were not differential during conidiation OR micafungin. This would indicate either (i) the BRE could have recombined into redundant, unused locations in the genome, or (ii) *brlA* could have roles in cellular process distinct from both conidiation and the response to cell-wall stress. Furthermore, there are 54 genes which have a BRE and that are only differential during conidiation. While our focus is the response to cell-wall stress, this list of 54 could be of great value to research groups focused solely on the asexual development cycle and physiology surrounding conidial development. Finally, the dynamic transcriptomic response of *A. nidulans* under synchronized conidiation is a dataset which has not been published before.

### Conclusions

BrlA is a transcription factor with a large dynamic range and many different roles. While well known to be a master regulator of conidiation, this work demonstrates that *brlA*, the *brlA* μORF, and *abaA* have divergent roles under cell-wall stress and during conidiation. The *brlA* μORF transcript is not differentially expressed during the conidiation response despite being differential under cell wall stress. *AbaA* is not differential during the cell-wall stress response, despite being upregulated 10-fold throughout conidiation. This work finds several transmembrane transporters that show altered expression levels in the absence of *brlA*. This work also demonstrates that the cell-wall stress response of *brlA* and the *brlA* μORF is mediated by MpkA. Finally, this work implies that there may be even more functionalities for BrlA based on the 226 genes with BREs which were not differential during either cell wall stress or conidiation. This work aligns with current literature on the subject and contributes to the mounting body of work that suggests *brlA* plays a role in cellular processes other than conidiation.

## Methods

### Plasmids, strains and media

Several *A. nidulans* strains were used in this work. *A. nidulans* A1405 was used in the dynamic transcriptomic response to micafungin. A4 was used in the dynamic transcriptomic response to synchronized conidiation. A1404 was the *mpkA* mutant strain used in the experiment shown in Figure 1. For the qPCR analysis, the wildtype strain was A1405 and the *brlA* mutant was A1826. YGV (0.5% Yeast Extract, 2% Glucose, with Vitamins and Hutner’s Trace Elements supplements)(*28, 29*) was used for all liquid media, and MAGV solid media (2% Glucose, 2% Malt Extract, 0.2% Peptone, 1.5% Agar, 0.95M Sucrose, with Vitamins and Hutner’s Trace Elements supplements)(*28, 29*) plates were used for all solid media, unless otherwise specified.

### Motif Search for Regulatory Motifs

The sequence of DNA which BrlA binds to, or the BrlA Response Element (BRE), is known (*27*), as is the whole genome sequence of *Aspergillus nidulans (30)*. We performed a genome-wide motif search for permutations of the semi-conserved BRE sequence: 5’- (C/A)(G/A)AGGG(G/A)-3’(*27*). We chose to search for the first six base pairs of this motif, both to reduce computational complexity and because 5’-CAAGGG-3’ is the most highly conserved BRE. The position and structure of the BrlA regulatory motif is mostly known because the upstream regions of genes known to be regulated by BrlA have been analyzed (*22, 27*). Based on previous studies, we use a sliding window size of 150 base pairs (bp). To allow for overlap and fuller coverage, the 150 bp sliding window is only shifted by 100 bp on each iteration of the search. Next, only regions beginning less than 1000 bp upstream from a start codon are considered. Considering these constraints, if three or more BREs appear within a window then that gene is saved as a possible BrlA target. This analysis yielded the 332 genes found in Supplementary File (S2).

### Transcriptomic data

The synchronized conidiation transcript data (Fig 1) were generated as follows. To generate sufficient mass of mycelium to harvest, an 8-liter carboy containing 4 liters of minimal medium media (1% Glucose, with Vitamins and Hutner’s Trace Elements supplements (*28, 29*) was inoculated with conidia to make a suspension with a final concentration of 10^6^ conidia/ml. Filter sterilized air was vigorously bubbled through the suspension to ensure aeration and agitation for 20 hr at room temperature, resulting in a logarithmic phase suspension of mycelial growth. Subsequently, 200 ml aliquots of the mycelial culture were harvested by vacuum filtration onto 9-cm Whatman no. 1 filter paper discs. Duplicate time point “0” samples was immediately frozen in liquid nitrogen. For the aerial time course, additional harvested mycelium on filter was placed onto 5 mm glass beads that were in 9-cm glass petri dishes containing liquid Aspergillus minimum medium to wick via the beads to maintain the moisture of the filter paper. Media levels were monitored and additional medium was added as needed to ensure filter papers remained wet. The time points were selected to ensure that key developmental stages were observed in a synchronous fashion (aerial hyphae ∼6 h, vesicle formation 6-12 h, metulae cell formation ∼12 h, phialide cell formation 12-18 h, conidiophore initiation 18-24 h, and mature conidiophore with conidial chains 24-48 h). Plates were numbered and placed in randomized positions under constant fluorescent light in a biosafety cabinet with a lowered glass sash and no airflow. Duplicate filter papers were collected for each time point. RNA was purified and cDNA constructed using the Illumina mRNA seq prep kit and sequenced on an Illumina GA II sequencer. This data will be housed on the Gene Expression Omnibus (GEO) GSE281842. The methods used to produce the transcriptomic time-course of A1405, and A1404, response to micafungin (Figure 2) has been published previously (*20, 21*), and the data is housed at GEO GSE136562. Briefly, spores were plated on MAGV, grown to maturation, harvested, and counted with hemocytometer such that 1 x 10^6^ fresh spores could be inoculated into a 250mL baffled flask with low pH (3.2) MAGV. This seed culture was grown for 12 hours (to germinate), and then used to inoculate a 2.5L Fernbach flask with 1.2L of MAGV. Fernbach flasks were treated with micafungin in the mid-exponential phase of growth. Samples were taken right before micafungin addition, and immediately after micafungin addition for the length of the time-course. Samples were liquid nitrogen frozen and crushed with mortar and pestle. RNA was extracted with Qiagen RNeasy RNA extraction kit, this was converted to cDNA with reverse transcriptase, and RNAseq data acquisition was performed at University of Nebraska Medical Center’s Bioinformatics and Systems Biology Core.

### qPCR

qPCR samples were taken from the culture flask during mid-exponential growth phase at the time of micafungin addition, and again one hour after. The culture flask work-up is the same procedure outlined above in the description of the micafungin treated *A. nidulans* transcriptomic data collection, and previously described in our lab (*20*). The procedure to design qPCR primers was adapted from the following work (*31*). qPCR primers were designed to have an amplicon length of 200 +/- 15 base pairs. Secondary and tertiary structures were checked using the methods described (*31*). Off target binding was avoided by using NCBI Blast and ensuring the amplicon was unique in the genome. The following pairs of primers (Supplementary File S3) are the qPCR primers used in the study, all of which were compared to the his2b protein for relative abundance. qPCR analysis was done via the relative fold change of starting concentration analysis, ΔΔN_0_, which incorporates qPCR efficiency (*32, 33*).

### DPoP

The Derivative Profiling omics Package is software which performs derivative profiling analysis on dynamic omics data to identify significantly differential signals from within the data set. A Pstat of 0.1 was to analyze the micafungin response data, and a Pstat of 0.15 was used for the synchronized conidiation transcriptomic response data. This increases confidence about what WAS differential to micafungin, and what WAS NOT differential to synchronized conidiation. Genes which DPoP found to be differential at any time point in the trajectory were kept for further analysis. This yielded 1839 genes differential to micafungin, and 2730 genes differential to synchronized conidiation (Supplementary File S2). These lists of genes, including those with a BRE, were then compared directly to create the Venn diagram seen in Figure 3.

## Acknowledgements

The authors would like to thank the NSF for their funding for this work. NSF Grant No. 2006189 supported researchers at UMBC, U. Conn, and Iowa State. NSF grant IOS-0716894 supported researchers at Texas A&M University

## Author contributions

HE was the primary researcher on the project responsible for aggregating/organizing the data, designing the confirmatory qPCR and for writing the manuscript. DE, HHW and BS produced the transcriptomic data set about *A. nidulans* under conditions of synchronized conidiation and wrote the accompanying methods section. JZ wrote the genome motif search algorithm to find BREs and the accompanying methods section. JL and KG aided with manuscript revisions and assisted in experimental design. HE did the DPoP analysis of the data sets and comparisons of the resulting lists. WH helped generate the transcriptomic data about the *mpka* mutant and helped develop the protocol to reproducibly grow the aconidial strain A1826 in shake flask. HE and MM executed the confirmatory qPCR experiments. AGD wrote the LinReg PCR analysis into an excel worksheet for the efficiency corrected qPCR analysis. BS, RS, SH, and MRM are co-PIs funding the work who provided guidance during research and revisions for the manuscript. MRM is the primary PI of the work and the corresponding author for this manuscript.

