## Supplementary Data S1 for "*Aspergillus nidulans* Transcription Factor BrlA is Utilized in a Conidiation-Independent Response to Cell-Wall Stress"

### Genome motif search

#### S1 – MATLAB Code to reproduce the genome motif search

```
function StartPOI = NidulansChromosomeSearch()
%% Info about this File:
% Written by: Joe Zavorskas
% Start: 9/9/2021 (Birthday Coding!)
% Last Edit: 10/4/2021

% This file will contain a workflow to search for clumps of brlA's
% experimentally proven binding motif (Chang; Timberlake, 1992) in the rest
% of the nidulans genome. This algorithm is an easy version of the motif
% finding problem. I will code the algorithm to generate all sequences and
% reverse compliments with up to "d" mutations of "CAAGGG" (given motif).
% I'll do this brute force for now, since I'm not sure if there is a more
% efficient method yet.

%% Input and manipulation of chromosome being searched.
% Goal is to get from many rows of character strings to a single char.
%%% I will add in functionality here to select 1 of the 8 chromosomes as
%%% well.
temp = regexp(fileread("Nidulans Chromo1.txt"), '\r?\n', 'split');
seq = vertcat(temp{1:end-1});
str = string(seq); singlestring = strjoin(str,"");
fullseq = char(singlestring);

%% Define target and generate the mutation neighborhood/reverse compliments.
% target(1) = {'CAAGGGG'};
% target(2) = {'AAAGGGG'};
% target(3) = {'CGAGGGG'};
% target(4) = {'AGAGGGG'};
% target(5) = {'CAAGGGA'};
% target(6) = {'AAAGGGA'};
% target(7) = {'CGAGGGA'};
% target(8) = {'AGAGGGA'};
target(1) = {'CAAGGG'};
target(2) = {'AAAGGG'};
target(3) = {'CGAGGG'};
target(4) = {'AGAGGG'};

% target(1) = {'CAAGGG'};
% target(2) = {'AAAGGGG'};
% target(3) = {'CGAGGGG'};
% target(4) = {'AGAGGGG'};
% target(5) = {'AAAGGGA'};
% target(6) = {'CGAGGGA'};
% target(7) = {'AGAGGGA'};

% d_NeighborsCell = Zavorskas_dNeighborhood(target,1);
```

```
ReverseComps = ReverseCompliments(target);

MatchCell = vertcat(target',ReverseComps);

%% Define window size and sampling rate for DNA sequences.

Window = 150;
% How far do we move the window before taking another sample?
SampleShift = 125;

% This hash table will store the start point of each window as its key. The
% value will be the number of times the target within one mutation or the
% reverse compliments appear in that window.
Counter = containers.Map('KeyType','double','ValueType','any');

%% For loop searching section

for WindowIter = 1:ceil(length(fullseq)/SampleShift)

    % Progress Updates

    if mod(WindowIter,10000) == 0

        Progress = WindowIter*SampleShift;
        format = 'Reached Position %d in the DNA.\n';
        fprintf(format,Progress)

    end

    TargetCount = 0;
    FirstValue = ((WindowIter-1)*SampleShift)+1;
    % Catch if statement to make sure the program doesn't run off the end
    % of the DNA strand with the given offset.
    try
        CheckSeq = fullseq(FirstValue:FirstValue+(Window-1));
    catch
        break
    end

    for Target = 1:length(MatchCell)

        WordCount(Target) = seqwordcount(CheckSeq,char(MatchCell(Target)));

    end

    TargetCount = sum(WordCount);

    % Map Update
    Counter(FirstValue) = TargetCount;

end

%% Analysis Section

StartPositions = keys(Counter);
```

```
Appearances = values(Counter);

StartPosMat = cell2mat(StartPositions)';
AppearMat = cell2mat(Appearances)';

MaxAppear = max(AppearMat)

OutputContainer = containers.Map('KeyType','double','ValueType','any');

for NumAppear = MaxAppear:-1:3

    idx = find(AppearMat == NumAppear);
    StartPOI = zeros(length(idx),1);

    for ID = 1:length(idx)

        StartPOI(ID) = StartPosMat(idx(ID));

    end

    OutputContainer(NumAppear) = StartPOI;

end

end
```

### Supplementary data

#### S2 – Lists of genes of interest from various analysis.

| Method of Analysis → | Contains BRE 332 Total | Differential to Micafungin 1839 Total | Differential during conidiation 2370 Total | Differential to micafungin, not dif. during conidiation 927 Total | Contains BRE, dif. to mica., not dif. during conidiation 29 Total |
|---|---|---|---|---|---|
| List of Significant Signals ↓ (Accession #) | AN3042 | AN7810 | AN1429 | AN7810 | AN5616 |
| | AN10085 | AN2393 | AN6426 | AN2393 | AN8330 |
| | AN1114 | AN1429 | AN0336 | AN2832 | AN7075 |
| | AN2860 | AN3581 | AN3446 | AN3581 | AN7836 |
| | AN10463 | AN9231 | AN0215 | AN9231 | AN0009 |
| | AN10892 | AN8159 | AN9216 | AN2817 | AN8595 |
| | AN1306 | AN2817 | AN9115 | AN8113 | AN5929 |
| | AN1176 | AN0336 | AN11188 | AN7983 | AN5311 |
| | AN11483 | AN0511 | AN8467 | AN8159 | AN1378 |
| | AN9280 | AN0215 | AN7261 | AN7622 | AN5385 |
| | AN4615 | AN0572 | AN5064 | AN6408 | AN0621 |
| | AN6848 | AN1261 | AN4740 | AN1651 | AN7541 |
| | AN4164 | AN7261 | AN11483 | AN0572 | AN0010 |
| | AN4984 | AN4885 | AN10699 | AN8650 | AN4975 |
| | AN9178 | AN8650 | AN11151 | AN2120 | AN5977 |
| | AN0906 | AN8986 | AN8176 | AN5037 | AN7828 |
| | AN4714 | AN5064 | AN11521 | AN4885 | AN2834 |
| | AN3554 | AN7073 | AN6022 | AN7112 | AN3553 |
| | AN10339 | AN2894 | AN0939 | AN4363 | AN6847 |
| | AN2704 | AN8107 | AN11571 | AN3523 | AN8155 |
| | AN4367 | AN6777 | AN7200 | AN8107 | AN6011 |
| | AN0077 | AN3402 | AN2704 | AN6236 | AN9280 |
| | AN7902 | AN6958 | AN5464 | AN3402 | AN3247 |
| | AN10853 | AN2559 | AN2699 | AN6777 | AN9165 |
| | AN5405 | AN6236 | AN2663 | AN2559 | AN3304 |
| | AN1188 | AN3783 | AN8647 | AN3783 | AN7580 |
| | AN0568 | AN3996 | AN7797 | AN9280 | AN8610 |
| | AN10431 | AN7327 | AN1198 | AN3996 | AN7860 |
| | AN6335 | AN6022 | AN1807 | AN7327 | AN3243 |
| | AN0017 | AN3780 | AN6542 | AN3780 | |
| | AN5371 | AN0374 | AN5834 | AN0374 | |
| | AN2284 | AN2704 | AN2032 | AN9337 | |
| | AN6694 | AN4970 | AN8654 | AN9374 | |
| | AN6725 | AN8477 | AN5335 | AN7780 | |
| | AN1638 | AN9374 | AN6392 | AN1315 | |
| | AN4233 | AN8647 | AN1584 | AN1066 | |
| | AN6517 | AN1315 | AN2789 | AN0036 | |
| | AN6814 | AN7797 | AN8363 | AN6657 | |
| | AN9272 | AN9313 | AN8507 | AN8411 | |
| | AN6743 | AN1066 | AN4605 | AN3276 | |
| | AN3304 | AN1198 | AN5542 | AN7827 | |
| | AN9201 | AN1807 | AN8944 | AN0915 | |
| | AN10870 | AN2099 | AN3267 | AN2542 | |
| | AN10230 | AN8411 | AN2888 | AN7097 | |
| | AN10430 | AN2032 | AN3305 | AN1829 | |
| | AN8595 | AN3276 | AN7356 | AN8429 | |
| | AN6391 | AN2346 | AN8390 | AN2025 | |
| | AN7418 | AN5335 | AN10605 | AN5893 | |
| | AN10232 | AN7097 | AN11101 | AN3889 | |

| | | | | |
|---|---|---|---|---|
| | AN3392<br>AN5095<br>AN0065<br>AN7828<br>AN0167<br>AN3093<br>AN8125<br>AN6255<br>AN10466<br>AN3411<br>AN4961<br>AN0166<br>AN6126<br>AN3143<br>AN4811<br>AN10644<br>AN0313<br>AN4657<br>AN3054<br>AN0264<br>AN10502<br>AN8513<br>AN8221<br>AN06230<br>AN2834<br>AN9177<br>AN10494<br>AN5688<br>AN3111<br>AN3949<br>AN7139<br>AN7252<br>AN10792<br>AN9208<br>AN4378<br>AN9165<br>AN0242<br>AN8330<br>AN5180<br>AN7560<br>AN0556<br>AN10799<br>AN4370<br>AN1844<br>AN5009<br>AN1378<br>AN2008<br>AN4975<br>AN1983<br>AN10833<br>AN10719<br>AN6305<br>AN5977<br>AN10656<br>AN8098<br>AN3798<br>AN7307<br>AN4136<br>AN4371<br>AN0265<br>AN2429<br>AN1560<br>AN11113<br>AN6683<br>AN8562<br>AN11082 | AN1829<br>AN8429<br>AN2025<br>AN2789<br>AN4640<br>AN8363<br>AN7417<br>AN1627<br>AN8659<br>AN2606<br>AN3162<br>AN1302<br>AN5312<br>AN4143<br>AN4605<br>AN5542<br>AN8944<br>AN3305<br>AN0565<br>AN8390<br>AN1257<br>AN5449<br>AN7877<br>AN7142<br>AN8124<br>AN7767<br>AN9503<br>AN2445<br>AN2028<br>AN8678<br>AN2557<br>AN5402<br>AN3503<br>AN2031<br>AN7800<br>AN9470<br>AN9443<br>AN7831<br>AN2810<br>AN3961<br>AN7794<br>AN0187<br>AN2378<br>AN5021<br>AN8731<br>AN1278<br>AN2943<br>AN5419<br>AN9163<br>AN4554<br>AN6096<br>AN7153<br>AN8463<br>AN7397<br>AN2891<br>AN2834<br>AN7528<br>AN0152<br>AN3201<br>AN5269<br>AN4852<br>AN8591<br>AN3195<br>AN1030<br>AN7409<br>AN6668 | AN9249<br>AN4938<br>AN6834<br>AN11611<br>AN4646<br>AN3058<br>AN3503<br>AN3326<br>AN7794<br>AN0187<br>AN2378<br>AN4313<br>AN9265<br>AN4292<br>AN2943<br>AN10259<br>AN4554<br>AN0967<br>AN5384<br>AN5227<br>AN6096<br>AN11091<br>AN1222<br>AN11035<br>AN6419<br>AN10368<br>AN2121<br>AN3201<br>AN0708<br>AN7887<br>AN9029<br>AN3228<br>AN4587<br>AN10741<br>AN9108<br>AN6668<br>AN11498<br>AN8378<br>AN2569<br>AN5193<br>AN10981<br>AN8959<br>AN1947<br>AN11157<br>AN10641<br>AN10862<br>AN7969<br>AN2535<br>AN0430<br>AN5138<br>AN11343<br>AN2193<br>AN1840<br>AN2337<br>AN3398<br>AN10656<br>AN9038<br>AN1231<br>AN0194<br>AN10977<br>AN7217<br>AN10782<br>AN9170<br>AN10997<br>AN5514<br>AN6667 | AN1627<br>AN4148<br>AN7417<br>AN8199<br>AN9012<br>AN2846<br>AN8659<br>AN2606<br>AN5312<br>AN8312<br>AN3162<br>AN3304<br>AN4143<br>AN8989<br>AN0565<br>AN8530<br>AN3327<br>AN8425<br>AN8595<br>AN5449<br>AN1257<br>AN8124<br>AN7621<br>AN2951<br>AN2028<br>AN2196<br>AN7226<br>AN8678<br>AN2557<br>AN7828<br>AN5402<br>AN9470<br>AN9443<br>AN8933<br>AN7831<br>AN8019<br>AN2384<br>AN8731<br>AN7383<br>AN5419<br>AN8365<br>AN0749<br>AN7875<br>AN8453<br>AN8463<br>AN7397<br>AN7079<br>AN2891<br>AN3132<br>AN2834<br>AN7528<br>AN0726<br>AN0152<br>AN5269<br>AN8446<br>AN8342<br>AN7962<br>AN2230<br>AN8540<br>AN6024<br>AN0723<br>AN7137<br>AN5393<br>AN9165<br>AN0983<br>AN7932 |

| | | | | |
|---|---|---|---|---|
| | AN0039<br>AN9101<br>AN9123<br>AN0856<br>AN2989<br>AN7242<br>AN1402<br>AN3958<br>AN7649<br>AN5741<br>AN9103<br>AN6246<br>AN9371<br>AN7580<br>AN8383<br>AN2229<br>AN0441<br>AN5643<br>AN5490<br>AN0009<br>AN6900<br>AN3784<br>AN4240<br>AN4001<br>AN5916<br>AN0010<br>AN7045<br>AN7884<br>AN10248<br>AN6334<br>AN8155<br>AN11440<br>AN1325<br>AN0154<br>AN9283<br>AN0986<br>AN9284<br>AN6224<br>AN7624<br>AN5989<br>AN2363<br>AN3377<br>AN10355<br>AN1955<br>AN9168<br>AN10864<br>AN08803<br>AN10283<br>AN4377<br>AN6284<br>AN6012<br>AN4936<br>AN7138<br>AN2885<br>AN11043<br>AN0532<br>AN5225<br>AN3243<br>AN4928<br>AN2897<br>AN3588<br>AN3797<br>AN6546<br>AN2946<br>AN5854<br>AN11031 | AN2230<br>AN4125<br>AN2569<br>AN6024<br>AN0723<br>AN7137<br>AN8959<br>AN0983<br>AN0615<br>AN8330<br>AN7531<br>AN6230<br>AN0788<br>AN7969<br>AN4660<br>AN2535<br>AN7282<br>AN9535<br>AN5977<br>AN2337<br>AN3988<br>AN9038<br>AN3398<br>AN8366<br>AN9335<br>AN6941<br>AN7119<br>AN9278<br>AN8620<br>AN2712<br>AN7217<br>AN5470<br>AN9487<br>AN9170<br>AN3194<br>AN0950<br>AN5537<br>AN8997<br>AN2034<br>AN2719<br>AN9259<br>AN8968<br>AN6667<br>AN5369<br>AN7868<br>AN2630<br>AN8137<br>AN5930<br>AN5129<br>AN4113<br>AN2565<br>AN7065<br>AN6021<br>AN8209<br>AN4619<br>AN7866<br>AN1035<br>AN5360<br>AN4337<br>AN8404<br>AN7791<br>AN6097<br>AN1857<br>AN7120<br>AN8106<br>AN8728 | AN4426<br>AN7868<br>AN0595<br>AN3264<br>AN5129<br>AN4113<br>AN5479<br>AN8209<br>AN5946<br>AN5360<br>AN4337<br>AN10649<br>AN6025<br>AN8728<br>AN8086<br>AN11452<br>AN6246<br>AN8153<br>AN7799<br>AN1915<br>AN3477<br>AN8251<br>AN0972<br>AN0697<br>AN2638<br>AN8976<br>AN9366<br>AN2044<br>AN10751<br>AN7960<br>AN11470<br>AN11471<br>AN7074<br>AN11492<br>AN10314<br>AN5130<br>AN1791<br>AN10044<br>AN11171<br>AN1049<br>AN11089<br>AN8915<br>AN3886<br>AN10199<br>AN3288<br>AN9339<br>AN6238<br>AN9284<br>AN7948<br>AN1576<br>AN7624<br>AN7992<br>AN8063<br>AN2401<br>AN7190<br>AN8484<br>AN11153<br>AN11034<br>AN3984<br>AN8547<br>AN7859<br>AN2573<br>AN3631<br>AN8909<br>AN11331<br>AN11362 | AN4330<br>AN3272<br>AN0615<br>AN8545<br>AN8330<br>AN6230<br>AN0788<br>AN4660<br>AN3570<br>AN0393<br>AN1378<br>AN7282<br>AN5437<br>AN8648<br>AN8999<br>AN9055<br>AN9296<br>AN4975<br>AN3573<br>AN1615<br>AN5977<br>AN6659<br>AN3988<br>AN2576<br>AN8685<br>AN8115<br>AN9335<br>AN7061<br>AN1144<br>AN7119<br>AN9278<br>AN8773<br>AN3109<br>AN2201<br>AN5470<br>AN0495<br>AN3194<br>AN9487<br>AN0950<br>AN5537<br>AN8997<br>AN2719<br>AN1185<br>AN8968<br>AN9046<br>AN5369<br>AN2630<br>AN8137<br>AN5218<br>AN2565<br>AN7065<br>AN6021<br>AN4619<br>AN1620<br>AN1035<br>AN9309<br>AN7791<br>AN8404<br>AN6097<br>AN1857<br>AN4815<br>AN8730<br>AN8105<br>AN2339<br>AN1590<br>AN5378 |

| | | | | |
|---|---|---|---|---|
| | AN9179 | AN8730 | AN2615 | AN0867 |
| | AN2288 | AN2339 | AN4299 | AN0786 |
| | AN2228 | AN1590 | AN4898 | AN8336 |
| | AN5887 | AN5546 | AN6641 | AN7196 |
| | AN0613 | AN0867 | AN4138 | AN3234 |
| | AN7348 | AN7053 | AN7811 | AN2468 |
| | AN0557 | AN0786 | AN1334 | AN3312 |
| | AN2968 | AN7799 | AN11446 | AN0009 |
| | AN4444 | AN8336 | AN9179 | AN8116 |
| | AN7716 | AN8251 | AN9020 | AN0028 |
| | AN4573 | AN0972 | AN11000 | AN2563 |
| | AN6291 | AN7196 | AN7893 | AN0010 |
| | AN8610 | AN3234 | AN7667 | AN7251 |
| | AN7860 | AN2638 | AN6427 | AN1838 |
| | AN1936 | AN3410 | AN1125 | AN1309 |
| | AN1567 | AN1580 | AN7881 | AN2778 |
| | AN2957 | AN3312 | AN6428 | AN2319 |
| | AN4985 | AN0009 | AN2548 | AN2381 |
| | AN4547 | AN7960 | AN7958 | AN6057 |
| | AN1715 | AN8116 | AN3873 | AN4106 |
| | AN5311 | AN0028 | AN2100 | AN0392 |
| | AN6590 | AN2563 | AN7769 | AN9479 |
| | AN2538 | AN0010 | AN3694 | AN9341 |
| | AN0855 | AN1914 | AN7213 | AN8727 |
| | AN1220 | AN5130 | AN5044 | AN8994 |
| | AN0253 | AN5564 | AN3611 | AN3763 |
| | AN3450 | AN1838 | AN5667 | AN8651 |
| | AN2437 | AN8241 | AN10152 | AN3782 |
| | AN0160 | AN2778 | AN9316 | AN3557 |
| | AN2967 | AN2381 | AN2729 | AN5477 |
| | AN8885 | AN0392 | AN4142 | AN9219 |
| | AN6187 | AN9479 | AN1603 | AN7272 |
| | AN5559 | AN8727 | AN8525 | AN6413 |
| | AN4960 | AN3782 | AN6421 | AN9524 |
| | AN2005 | AN3886 | AN0479 | AN7488 |
| | AN4366 | AN3763 | AN11610 | AN9013 |
| | AN0815 | AN8994 | AN2372 | AN1199 |
| | AN1689 | AN2804 | AN6181 | AN5024 |
| | AN6847 | AN9339 | AN4137 | AN0644 |
| | AN3363 | AN9315 | AN1551 | AN7110 |
| | AN2741 | AN9284 | AN9510 | AN1622 |
| | AN4803 | AN7272 | AN4649 | AN6131 |
| | AN9471 | AN6413 | AN3993 | AN0645 |
| | AN5617 | AN9524 | AN10896 | AN3997 |
| | AN1635 | AN5024 | AN8658 | AN6315 |
| | AN7676 | AN1576 | AN11647 | AN0860 |
| | AN10404 | AN1199 | AN3609 | AN6153 |
| | AN10220 | AN7992 | AN3339 | AN9045 |
| | AN6221 | AN7110 | AN4366 | AN8451 |
| | AN2236 | AN1622 | AN8529 | AN8479 |
| | AN7650 | AN6131 | AN10027 | AN0231 |
| | AN5385 | AN3284 | AN2931 | AN8008 |
| | AN6257 | AN0645 | AN10421 | AN3987 |
| | AN7231 | AN3997 | AN3237 | AN2698 |
| | AN5534 | AN0860 | AN8409 | AN7972 |
| | AN9389 | AN9045 | AN10086 | AN6535 |
| | AN6263 | AN8484 | AN2617 | AN9277 |
| | AN10447 | AN3984 | AN1181 | AN1686 |
| | AN6602 | AN8547 | AN7681 | AN0732 |
| | AN11174 | AN1837 | AN8046 | AN8460 |
| | AN10413 | AN7859 | AN5957 | AN4924 |
| | AN6552 | AN2573 | AN1041 | AN3243 |
| | AN6518 | AN7322 | AN6406 | AN2826 |
| | AN1745 | AN4480 | AN3887 | AN8436 |
| | AN1998 | AN8909 | AN4139 | AN7592 |
| | AN5853 | AN1686 | AN6661 | AN2325 |

| | | | | |
|---|---|---|---|---|
| | AN0409 | AN0732 | AN9030 | AN7669 |
| | AN2321 | AN2344 | AN0478 | AN5976 |
| | AN5616 | AN8436 | AN2682 | AN5091 |
| | AN11320 | AN7811 | AN6660 | AN7855 |
| | AN10839 | AN9026 | AN5602 | AN0899 |
| | AN7836 | AN3077 | AN0322 | AN6576 |
| | AN0868 | AN7669 | AN5660 | AN3585 |
| | AN10899 | AN9179 | AN3261 | AN1570 |
| | AN5929 | AN5091 | AN6378 | AN5841 |
| | AN6898 | AN7893 | AN8464 | AN5690 |
| | AN6676 | AN7855 | AN8550 | AN1581 |
| | AN1999 | AN3585 | AN6933 | AN3248 |
| | AN5653 | AN1570 | AN7697 | AN0400 |
| | AN2132 | AN5690 | AN8310 | AN5290 |
| | AN1935 | AN1581 | AN3357 | AN3214 |
| | AN3218 | AN1125 | AN6026 | AN0364 |
| | AN5151 | AN7881 | AN5339 | AN8445 |
| | AN2348 | AN5290 | AN2368 | AN7891 |
| | AN10869 | AN8445 | AN4991 | AN3561 |
| | AN1827 | AN7891 | AN8627 | AN2781 |
| | AN1269 | AN3561 | AN2012 | AN8607 |
| | AN7075 | AN2781 | AN9131 | AN7069 |
| | AN10604 | AN7850 | AN9352 | AN9381 |
| | AN5695 | AN8607 | AN7768 | AN0011 |
| | AN11590 | AN7069 | AN6098 | AN9314 |
| | AN5255 | AN3873 | AN5543 | AN3200 |
| | AN10177 | AN9381 | AN5743 | AN0482 |
| | AN5630 | AN0011 | AN2421 | AN8405 |
| | AN6516 | AN7769 | AN11114 | AN9226 |
| | AN6759 | AN0482 | AN5228 | AN4215 |
| | AN4328 | AN7213 | AN6256 | AN8422 |
| | AN6538 | AN9226 | AN4180 | AN8279 |
| | AN2958 | AN4215 | AN8335 | AN7860 |
| | AN11404 | AN5044 | AN10477 | AN0548 |
| | AN0314 | AN3611 | AN2877 | AN0612 |
| | AN5163 | AN8279 | AN2035 | AN7812 |
| | AN11383 | AN7860 | AN7377 | AN0169 |
| | AN1672 | AN8405 | AN5511 | AN6168 |
| | AN0979 | AN9316 | AN10778 | AN7558 |
| | AN9390 | AN6784 | AN2621 | AN5397 |
| | AN3110 | AN2729 | AN5654 | AN0332 |
| | AN2006 | AN0612 | AN5846 | AN0688 |
| | AN10094 | AN4142 | AN7021 | AN8583 |
| | AN3247 | AN7812 | AN6060 | AN1092 |
| | AN4332 | AN8353 | AN0378 | AN3967 |
| | AN7883 | AN6168 | AN11485 | AN8509 |
| | AN2282 | AN7558 | AN8579 | AN7933 |
| | AN6750 | AN8583 | AN4734 | AN2360 |
| | AN5026 | AN5397 | AN6811 | AN5478 |
| | AN7995 | AN0688 | AN7333 | AN7233 |
| | AN2953 | AN1092 | AN7614 | AN1744 |
| | AN8774 | AN8509 | AN4131 | AN2200 |
| | AN3148 | AN2372 | AN0387 | AN8612 |
| | AN0951 | AN1542 | AN11092 | AN8611 |
| | AN0223 | AN7933 | AN5833 | AN4257 |
| | AN10630 | AN2360 | AN3390 | AN8781 |
| | AN7541 | AN2599 | AN8140 | AN4769 |
| | AN3020 | AN5478 | AN9263 | AN5046 |
| | AN6285 | AN7233 | AN11650 | AN7022 |
| | AN10428 | AN5310 | AN4121 | AN5651 |
| | AN10557 | AN1744 | AN6235 | AN0015 |
| | AN6321 | AN5859 | AN1898 | AN6949 |
| | AN11090 | AN2225 | AN7843 | AN0148 |
| | AN4802 | AN8349 | AN8318 | AN0964 |
| | AN11539 | AN8333 | AN7160 | AN9210 |
| | AN6011 | AN8011 | AN2677 | AN8262 |

| | | | | |
|---|---|---|---|---|
| | AN0835 | AN8781 | AN4650 | AN2361 |
| | AN1783 | AN8135 | AN2526 | AN5354 |
| | AN2884 | AN5046 | AN4588 | AN7949 |
| | AN0621 | AN6949 | AN5496 | AN5408 |
| | AN10464 | AN0964 | AN7386 | AN2187 |
| | AN11095 | AN0148 | AN11085 | AN5935 |
| | AN6836 | AN3339 | AN2427 | AN5918 |
| | AN5368 | AN9210 | AN2475 | AN4485 |
| | AN3553 | AN1803 | AN2502 | AN6847 |
| | AN5583 | AN2361 | AN7516 | AN2842 |
| | AN5356 | AN2544 | AN4592 | AN5427 |
| | AN0902 | AN5408 | AN2678 | AN5320 |
| | AN2124 | AN2187 | AN7201 | AN2027 |
| | AN0346 | AN5918 | AN3068 | AN8093 |
| | AN2937 | AN3237 | AN5914 | AN3241 |
| | AN9517 | AN8409 | AN4252 | AN8693 |
| | AN5845 | AN2842 | AN5497 | AN5286 |
| | AN0408 | AN5320 | AN10565 | AN1184 |
| | AN5182 | AN2027 | AN11605 | AN2192 |
| | | AN8046 | AN5317 | AN7408 |
| | | AN3241 | AN7118 | AN3951 |
| | | AN8693 | AN8614 | AN7270 |
| | | AN1184 | AN4677 | AN9006 |
| | | AN5467 | AN8337 | AN5089 |
| | | AN7270 | AN8412 | AN2583 |
| | | AN9006 | AN10289 | AN4151 |
| | | AN6406 | AN9174 | AN0941 |
| | | AN3887 | AN10207 | AN5385 |
| | | AN6661 | AN5325 | AN7095 |
| | | AN4151 | AN5900 | AN7410 |
| | | AN5385 | AN2380 | AN2572 |
| | | AN7410 | AN5086 | AN3665 |
| | | AN7402 | AN6318 | AN7515 |
| | | AN4325 | AN9053 | AN5029 |
| | | AN2682 | AN7598 | AN8538 |
| | | AN6660 | AN8920 | AN2805 |
| | | AN3665 | AN10984 | AN4156 |
| | | AN5602 | AN8508 | AN3344 |
| | | AN0322 | AN0710 | AN8754 |
| | | AN2622 | AN3042 | AN2412 |
| | | AN3261 | AN6754 | AN3384 |
| | | AN7515 | AN10789 | AN9295 |
| | | AN6378 | AN3283 | AN1887 |
| | | AN8550 | AN2768 | AN9056 |
| | | AN2805 | AN2746 | AN1318 |
| | | AN4156 | AN10728 | AN3331 |
| | | AN6377 | AN3245 | AN4115 |
| | | AN8754 | AN0301 | AN0488 |
| | | AN9295 | AN7839 | AN9282 |
| | | AN5339 | AN1788 | AN5616 |
| | | AN3331 | AN8906 | AN8993 |
| | | AN8352 | AN2470 | AN3321 |
| | | AN2097 | AN5260 | AN6541 |
| | | AN8627 | AN5746 | AN7836 |
| | | AN4115 | AN7857 | AN8449 |
| | | AN9282 | AN9261 | AN5964 |
| | | AN9352 | AN2836 | AN2869 |
| | | AN8993 | AN7752 | AN5929 |
| | | AN8621 | AN7984 | AN8642 |
| | | AN6541 | AN8761 | AN2655 |
| | | AN6098 | AN5190 | AN2707 |
| | | AN5942 | AN6926 | AN5975 |
| | | AN5228 | AN5818 | AN2351 |
| | | AN2707 | AN6443 | AN4909 |
| | | AN2655 | AN1997 | AN2671 |
| | | AN2351 | AN1017 | AN7841 |

| | | | | |
|---|---|---|---|---|
| | | AN2035<br>AN2671<br>AN5511<br>AN6955<br>AN2823<br>AN1321<br>AN2621<br>AN8277<br>AN3892<br>AN5654<br>AN5846<br>AN7021<br>AN2669<br>AN0300<br>AN1747<br>AN3963<br>AN1613<br>AN0378<br>AN8399<br>AN5259<br>AN7668<br>AN5079<br>AN7333<br>AN8551<br>AN8424<br>AN1288<br>AN3151<br>AN2347<br>AN4131<br>AN2672<br>AN2186<br>AN2680<br>AN6118<br>AN7020<br>AN3390<br>AN9240<br>AN8140<br>AN9263<br>AN0992<br>AN3247<br>AN8593<br>AN7876<br>AN6235<br>AN1323<br>AN0019<br>AN8186<br>AN1898<br>AN7843<br>AN8893<br>AN9372<br>AN0032<br>AN7160<br>AN8548<br>AN2642<br>AN1548<br>AN2677<br>AN2706<br>AN3859<br>AN1600<br>AN7541<br>AN9161<br>AN9247<br>AN7266<br>AN7412<br>AN1901<br>AN7386 | AN7466<br>AN8637<br>AN6930<br>AN11555<br>AN8265<br>AN8315<br>AN2923<br>AN2640<br>AN8995<br>AN6543<br>AN11508<br>AN6649<br>AN0610<br>AN11488<br>AN7639<br>AN2721<br>AN8589<br>AN3697<br>AN6830<br>AN7126<br>AN1969<br>AN2394<br>AN11425<br>AN1893<br>AN3774<br>AN3392<br>AN1896<br>AN9051<br>AN3679<br>AN8741<br>AN2038<br>AN10344<br>AN3790<br>AN10099<br>AN8521<br>AN4127<br>AN2818<br>AN9087<br>AN7178<br>AN7380<br>AN2170<br>AN5993<br>AN3567<br>AN0353<br>AN11403<br>AN3205<br>AN6018<br>AN8922<br>AN0524<br>AN11486<br>AN9388<br>AN9253<br>AN7234<br>AN5780<br>AN9040<br>AN3703<br>AN4392<br>AN3949<br>AN1169<br>AN6856<br>AN3358<br>AN2218<br>AN1566<br>AN7379<br>AN2942<br>AN0801 | AN3238<br>AN0456<br>AN8302<br>AN8277<br>AN3892<br>AN6509<br>AN0660<br>AN0300<br>AN5019<br>AN3963<br>AN1613<br>AN1747<br>AN7075<br>AN9301<br>AN0208<br>AN5259<br>AN7668<br>AN8551<br>AN1288<br>AN7549<br>AN8341<br>AN1692<br>AN7263<br>AN2186<br>AN2672<br>AN2040<br>AN2680<br>AN7900<br>AN6118<br>AN7020<br>AN1556<br>AN6944<br>AN9240<br>AN8408<br>AN1631<br>AN3247<br>AN0992<br>AN0019<br>AN1323<br>AN8186<br>AN5226<br>AN7063<br>AN7018<br>AN8893<br>AN7532<br>AN8632<br>AN0014<br>AN2642<br>AN8548<br>AN7818<br>AN1548<br>AN9372<br>AN2800<br>AN2706<br>AN3859<br>AN7392<br>AN1600<br>AN7541<br>AN8348<br>AN7266<br>AN1901<br>AN7264<br>AN2941<br>AN7161<br>AN0798<br>AN5045 |

| | | | | |
|---|---|---|---|---|
| | | AN7516<br>AN2941<br>AN7161<br>AN5045<br>AN2727<br>AN2678<br>AN1609<br>AN8487<br>AN0619<br>AN8465<br>AN8334<br>AN8006<br>AN7484<br>AN8585<br>AN8948<br>AN9318<br>AN0621<br>AN7385<br>AN7118<br>AN2469<br>AN6023<br>AN8337<br>AN2792<br>AN7634<br>AN5056<br>AN8412<br>AN0074<br>AN9174<br>AN8101<br>AN8543<br>AN7946<br>AN9053<br>AN5086<br>AN7806<br>AN0833<br>AN8920<br>AN9223<br>AN3571<br>AN8508<br>AN5845<br>AN0971<br>AN7094<br>AN8060<br>AN3391<br>AN4019<br>AN8934<br>AN0402<br>AN3283<br>AN2746<br>AN3615<br>AN3280<br>AN2387<br>AN9321<br>AN3353<br>AN6810<br>AN2470<br>AN0220<br>AN4905<br>AN9261<br>AN3511<br>AN3303<br>AN8641<br>AN6237<br>AN4607<br>AN7984<br>AN6672 | AN1047<br>AN8392<br>AN8468<br>AN10838<br>AN0639<br>AN6461<br>AN6890<br>AN3550<br>AN11564<br>AN1642<br>AN11642<br>AN8329<br>AN4135<br>AN4833<br>AN11413<br>AN3549<br>AN6782<br>AN7781<br>AN1029<br>AN0146<br>AN2119<br>AN3536<br>AN4424<br>AN5338<br>AN7084<br>AN6703<br>AN8556<br>AN8139<br>AN9088<br>AN8967<br>AN1043<br>AN5634<br>AN6943<br>AN2042<br>AN0493<br>AN3481<br>AN6471<br>AN8502<br>AN10665<br>AN0857<br>AN3226<br>AN8998<br>AN8068<br>AN2855<br>AN0216<br>AN3265<br>AN8625<br>AN0147<br>AN0473<br>AN2613<br>AN7990<br>AN0051<br>AN9441<br>AN2676<br>AN1801<br>AN9283<br>AN2525<br>AN3613<br>AN11460<br>AN2913<br>AN0429<br>AN10148<br>AN6954<br>AN8325<br>AN11279<br>AN11646 | AN4173<br>AN2727<br>AN1609<br>AN9128<br>AN8692<br>AN8487<br>AN8334<br>AN1928<br>AN5213<br>AN7484<br>AN8585<br>AN8948<br>AN7518<br>AN0621<br>AN7854<br>AN0546<br>AN2469<br>AN6023<br>AN7634<br>AN7882<br>AN3553<br>AN5665<br>AN0074<br>AN9094<br>AN8543<br>AN2796<br>AN2127<br>AN3139<br>AN7946<br>AN2624<br>AN7806<br>AN2403<br>AN6942<br>AN0833<br>AN9223<br>AN7452<br>AN3571<br>AN7054<br>AN0971<br>AN7094<br>AN8060<br>AN4019<br>AN8934<br>AN0402<br>AN7823<br>AN3615<br>AN3289<br>AN8778<br>AN8582<br>AN0737<br>AN3280<br>AN5093<br>AN2387<br>AN6359<br>AN3858<br>AN3353<br>AN5276<br>AN7635<br>AN4905<br>AN3255<br>AN3511<br>AN8552<br>AN3303<br>AN0531<br>AN6237<br>AN7176 |

| | | | | |
|---|---|---|---|---|
| | | AN0933<br>AN1997<br>AN8637<br>AN6930<br>AN6585<br>AN7965<br>AN8265<br>AN1855<br>AN6957<br>AN3474<br>AN7317<br>AN3998<br>AN2640<br>AN8995<br>AN8102<br>AN4108<br>AN5843<br>AN6649<br>AN9175<br>AN0610<br>AN6621<br>AN2574<br>AN0617<br>AN0760<br>AN8589<br>AN0234<br>AN6830<br>AN8561<br>AN1311<br>AN1969<br>AN0230<br>AN1893<br>AN4774<br>AN8308<br>AN0683<br>AN9002<br>AN1897<br>AN7418<br>AN1896<br>AN6869<br>AN8741<br>AN3508<br>AN2038<br>AN8340<br>AN2407<br>AN3991<br>AN8521<br>AN2036<br>AN6691<br>AN4127<br>AN9181<br>AN5944<br>AN0468<br>AN2382<br>AN0516<br>AN7178<br>AN7380<br>AN7956<br>AN3872<br>AN2588<br>AN2170<br>AN8351<br>AN1573<br>AN3299<br>AN6525<br>AN9502 | AN1088<br>AN6882<br>AN3197<br>AN4107<br>AN8656<br>AN8680<br>AN2881<br>AN2309<br>AN6482<br>AN10626<br>AN7905<br>AN7557<br>AN0122<br>AN7927<br>AN9288<br>AN6172<br>AN11374<br>AN10996<br>AN6773<br>AN9250<br>AN7826<br>AN1138<br>AN10256<br>AN10730<br>AN2532<br>AN2602<br>AN3085<br>AN6820<br>AN2984<br>AN4972<br>AN5176<br>AN11652<br>AN5294<br>AN8044<br>AN10993<br>AN7519<br>AN2455<br>AN1681<br>AN8802<br>AN4727<br>AN10573<br>AN10183<br>AN10343<br>AN8609<br>AN2783<br>AN2359<br>AN0235<br>AN9194<br>AN10210<br>AN10321<br>AN5756<br>AN7067<br>AN8523<br>AN0530<br>AN1034<br>AN6154<br>AN10258<br>AN6946<br>AN8202<br>AN2238<br>AN2665<br>AN11544<br>AN0489<br>AN7520<br>AN0318<br>AN1831 | AN5370<br>AN4607<br>AN1623<br>AN6672<br>AN0933<br>AN1574<br>AN9303<br>AN3476<br>AN7070<br>AN7867<br>AN0016<br>AN1664<br>AN9351<br>AN6936<br>AN7965<br>AN1322<br>AN6585<br>AN1855<br>AN7391<br>AN6751<br>AN6957<br>AN3474<br>AN7910<br>AN4108<br>AN2950<br>AN9175<br>AN6621<br>AN2574<br>AN0617<br>AN5507<br>AN0234<br>AN0760<br>AN8561<br>AN7813<br>AN2679<br>AN9332<br>AN8308<br>AN4774<br>AN4854<br>AN0683<br>AN7842<br>AN6671<br>AN6959<br>AN6869<br>AN0958<br>AN2550<br>AN3531<br>AN5053<br>AN3508<br>AN9044<br>AN8340<br>AN2407<br>AN8955<br>AN3991<br>AN6691<br>AN9181<br>AN2036<br>AN3396<br>AN8160<br>AN2340<br>AN2198<br>AN1882<br>AN2382<br>AN0516<br>AN1573<br>AN5068 |

| | | | | |
|---|---|---|---|---|
| | | AN3978<br>AN6144<br>AN6018<br>AN3888<br>AN1509<br>AN8922<br>AN0524<br>AN4109<br>AN7931<br>AN8638<br>AN8513<br>AN6636<br>AN0944<br>AN1841<br>AN8371<br>AN2724<br>AN9207<br>AN3202<br>AN5030<br>AN4608<br>AN7379<br>AN0499<br>AN7413<br>AN6239<br>AN8392<br>AN1604<br>AN0639<br>AN7837<br>AN3901<br>AN8329<br>AN7393<br>AN8890<br>AN4135<br>AN3549<br>AN5541<br>AN4081<br>AN7781<br>AN1029<br>AN3536<br>AN7084<br>AN0609<br>AN8594<br>AN5338<br>AN8556<br>AN2593<br>AN8139<br>AN8953<br>AN8132<br>AN9088<br>AN1723<br>AN8527<br>AN6943<br>AN0493<br>AN9014<br>AN6935<br>AN9185<br>AN4144<br>AN2531<br>AN8998<br>AN3259<br>AN8068<br>AN0935<br>AN7580<br>AN3512<br>AN7068<br>AN9180 | AN6772<br>AN6379<br>AN8380<br>AN9015<br>AN4059<br>AN3409<br>AN7838<br>AN4154<br>AN8510<br>AN4773<br>AN2055<br>AN9401<br>AN8377<br>AN7275<br>AN2248<br>AN6148<br>AN10288<br>AN4157<br>AN5329<br>AN7539<br>AN7782<br>AN5624<br>AN1650<br>AN4400<br>AN8951<br>AN8379<br>AN5587<br>AN5313<br>AN8339<br>AN0550<br>AN7903<br>AN0759<br>AN7244<br>AN9248<br>AN3586<br>AN10457<br>AN10632<br>AN11207<br>AN5032<br>AN5551<br>AN3218<br>AN10869<br>AN3175<br>AN9184<br>AN5050<br>AN7116<br>AN3612<br>AN0751<br>AN6425<br>AN2607<br>AN3482<br>AN2605<br>AN4049<br>AN10261<br>AN10386<br>AN3515<br>AN10937<br>AN0513<br>AN2051<br>AN2549<br>AN5245<br>AN9252<br>AN11583<br>AN4586<br>AN6273<br>AN1036 | AN3299<br>AN5938<br>AN4627<br>AN2675<br>AN9502<br>AN3978<br>AN3420<br>AN6144<br>AN5417<br>AN1509<br>AN3888<br>AN2320<br>AN7931<br>AN4109<br>AN0944<br>AN0946<br>AN2724<br>AN9207<br>AN8962<br>AN8514<br>AN4711<br>AN3202<br>AN5030<br>AN9220<br>AN4608<br>AN6239<br>AN5937<br>AN1582<br>AN7961<br>AN7281<br>AN1644<br>AN7837<br>AN5831<br>AN2004<br>AN7393<br>AN9206<br>AN8617<br>AN7082<br>AN4081<br>AN5110<br>AN5541<br>AN8630<br>AN8594<br>AN0609<br>AN0158<br>AN3406<br>AN2593<br>AN8953<br>AN1723<br>AN8527<br>AN3713<br>AN7879<br>AN2039<br>AN8779<br>AN1630<br>AN6873<br>AN9185<br>AN2531<br>AN6718<br>AN1835<br>AN5664<br>AN0935<br>AN3273<br>AN0509<br>AN0750<br>AN4807 |

| | | | | |
|---|---|---|---|---|
| | | AN5316 | AN6951 | AN7580 |
| | | AN0216 | AN2821 | AN3512 |
| | | AN3265 | AN4567 | AN7916 |
| | | AN2609 | AN7323 | AN7068 |
| | | AN5556 | AN5480 | AN4195 |
| | | AN2708 | AN8814 | AN0601 |
| | | AN0551 | AN0467 | AN8155 |
| | | AN6961 | AN10491 | AN7267 |
| | | AN2625 | AN2585 | AN0024 |
| | | AN2613 | AN5326 | AN2708 |
| | | AN7990 | AN8220 | AN9139 |
| | | AN7384 | AN4859 | AN7647 |
| | | AN8480 | AN6386 | AN1450 |
| | | AN0563 | AN3101 | AN0551 |
| | | AN2676 | AN7637 | AN1742 |
| | | AN0918 | AN1645 | AN6961 |
| | | AN8549 | AN3973 | AN1591 |
| | | AN2787 | AN11090 | AN7152 |
| | | AN4908 | AN4606 | AN0563 |
| | | AN9190 | AN9246 | AN0918 |
| | | AN3613 | AN0652 | AN1903 |
| | | AN0151 | AN10761 | AN2814 |
| | | AN0542 | AN5061 | AN8549 |
| | | AN2913 | AN11527 | AN2787 |
| | | AN2788 | AN5319 | AN3475 |
| | | AN0404 | AN7663 | AN4908 |
| | | AN6954 | AN2221 | AN9190 |
| | | AN0523 | AN4582 | AN3252 |
| | | AN4950 | AN5349 | AN0151 |
| | | AN6882 | AN0510 | AN0542 |
| | | AN2568 | AN7401 | AN2788 |
| | | AN4141 | AN9031 | AN0404 |
| | | AN7833 | AN1473 | AN9427 |
| | | AN4107 | AN8151 | AN8433 |
| | | AN8930 | AN3562 | AN4950 |
| | | AN0157 | AN0859 | AN4128 |
| | | AN8662 | AN2610 | AN7833 |
| | | AN8913 | AN7165 | AN1203 |
| | | AN5016 | AN11333 | AN6809 |
| | | AN9245 | AN8949 | AN8930 |
| | | AN2286 | AN9191 | AN9121 |
| | | AN7199 | AN10582 | AN0157 |
| | | AN4105 | AN10141 | AN8662 |
| | | AN1571 | AN8426 | AN4357 |
| | | AN7083 | AN1067 | AN8338 |
| | | AN0517 | AN10279 | AN5016 |
| | | AN7557 | AN10733 | AN9245 |
| | | AN8907 | AN0638 | AN4146 |
| | | AN1541 | AN4213 | AN7199 |
| | | AN7081 | AN0100 | AN4105 |
| | | AN8904 | AN11345 | AN7083 |
| | | AN7826 | AN9385 | AN0517 |
| | | AN8555 | AN6317 | AN3496 |
| | | AN8910 | AN1937 | AN2330 |
| | | AN0022 | AN3862 | AN0515 |
| | | AN2471 | AN6135 | AN2199 |
| | | AN8374 | AN3893 | AN9456 |
| | | AN7772 | AN8996 | AN2310 |
| | | AN7319 | AN1900 | AN7089 |
| | | AN2532 | AN3052 | AN1541 |
| | | AN8084 | AN7793 | AN3494 |
| | | AN2696 | AN4615 | AN2197 |
| | | AN5301 | AN1859 | AN3253 |
| | | AN0338 | AN5652 | AN8904 |
| | | AN1575 | AN5943 | AN7896 |
| | | AN8515 | AN0779 | AN7577 |

| | | | | |
|---|---|---|---|---|
| | | AN5271 | AN7204 | AN3282 |
| | | AN0498 | AN10366 | AN7011 |
| | | AN8018 | AN10891 | AN2795 |
| | | AN7944 | AN3227 | AN2827 |
| | | AN3393 | AN11514 | AN3479 |
| | | AN8323 | AN11346 | AN7772 |
| | | AN5311 | AN0335 | AN7319 |
| | | AN5294 | AN10909 | AN7191 |
| | | AN6129 | AN2901 | AN8084 |
| | | AN3249 | AN8175 | AN5301 |
| | | AN9212 | AN2589 | AN0338 |
| | | AN7519 | AN11295 | AN5509 |
| | | AN8360 | AN1717 | AN8515 |
| | | AN2455 | AN1143 | AN1575 |
| | | AN1681 | AN8808 | AN8610 |
| | | AN5502 | AN1240 | AN5271 |
| | | AN8802 | AN8558 | AN0498 |
| | | AN3849 | AN5364 | AN7944 |
| | | AN6624 | AN7959 | AN7398 |
| | | AN2608 | AN1214 | AN2694 |
| | | AN8085 | AN11308 | AN8108 |
| | | AN3983 | AN2883 | AN8927 |
| | | AN4152 | AN8570 | AN3023 |
| | | AN1451 | AN2530 | AN3393 |
| | | AN8616 | AN2564 | AN6487 |
| | | AN8609 | AN11441 | AN9200 |
| | | AN2783 | AN9054 | AN8323 |
| | | AN2359 | AN11047 | AN0600 |
| | | AN3488 | AN1652 | AN5311 |
| | | AN9036 | AN3292 | AN5072 |
| | | AN9194 | AN4029 | AN6129 |
| | | AN4111 | AN11059 | AN3545 |
| | | AN3275 | AN4051 | AN1594 |
| | | AN0512 | AN2813 | AN3249 |
| | | AN7067 | AN6665 | AN9242 |
| | | AN3347 | AN5281 | AN9212 |
| | | AN1852 | AN8398 | AN4474 |
| | | AN6281 | AN8639 | AN8158 |
| | | AN0530 | AN3884 | AN7268 |
| | | AN1752 | AN7470 | AN8360 |
| | | AN2841 | AN1616 | AN7702 |
| | | AN1034 | AN4132 | AN5502 |
| | | AN6154 | AN11160 | AN7600 |
| | | AN6946 | AN9189 | AN3849 |
| | | AN0553 | AN7395 | AN2608 |
| | | AN0285 | AN11016 | AN8085 |
| | | AN5394 | AN10040 | AN3983 |
| | | AN5611 | AN6027 | AN1451 |
| | | AN8452 | AN2835 | AN3488 |
| | | AN0489 | AN1107 | AN8616 |
| | | AN0735 | AN2203 | AN9001 |
| | | AN6772 | AN10050 | AN3275 |
| | | AN1831 | AN8592 | AN9375 |
| | | AN8222 | AN8149 | AN0512 |
| | | AN4120 | AN2191 | AN6662 |
| | | AN2110 | AN4402 | AN3347 |
| | | AN7838 | AN9177 | AN6281 |
| | | AN5383 | AN2864 | AN8143 |
| | | AN4372 | AN11349 | AN0622 |
| | | AN8979 | AN2547 | AN1752 |
| | | AN8622 | AN6319 | AN2841 |
| | | AN8510 | AN1263 | AN4460 |
| | | AN9224 | AN9198 | AN2892 |
| | | AN8377 | AN0528 | AN2649 |
| | | AN3776 | AN4644 | AN5396 |
| | | AN7275 | AN2755 | AN1938 |

| | | | | |
|---|---|---|---|---|
| | | AN7202 | AN3233 | AN0553 |
| | | AN0366 | AN6881 | AN0285 |
| | | AN4157 | AN9531 | AN5555 |
| | | AN5329 | AN0744 | AN5394 |
| | | AN1839 | AN7939 | AN0635 |
| | | AN7539 | AN2653 | AN8596 |
| | | AN5280 | AN8732 | AN3387 |
| | | AN1215 | AN2061 | AN8452 |
| | | AN7098 | AN3207 | AN6739 |
| | | AN5240 | AN7411 | AN5076 |
| | | AN1140 | AN3078 | AN0735 |
| | | AN2509 | AN8640 | AN7066 |
| | | AN8458 | AN0891 | AN4117 |
| | | AN1677 | AN7181 | AN8222 |
| | | AN9476 | AN3196 | AN2714 |
| | | AN3891 | AN9495 | AN2110 |
| | | AN8397 | AN11113 | AN5383 |
| | | AN0550 | AN10807 | AN4372 |
| | | AN2811 | AN4124 | AN1073 |
| | | AN2352 | AN0403 | AN7814 |
| | | AN0365 | AN2562 | AN9224 |
| | | AN7390 | AN8563 | AN8935 |
| | | AN0759 | AN9262 | AN3271 |
| | | AN6966 | AN7147 | AN4879 |
| | | AN8666 | AN8250 | AN3776 |
| | | AN2357 | AN10692 | AN7057 |
| | | AN6967 | AN2224 | AN0366 |
| | | AN9248 | AN11520 | AN1839 |
| | | AN3533 | AN1402 | AN5034 |
| | | AN1008 | AN1529 | AN5280 |
| | | AN5355 | AN3314 | AN7080 |
| | | AN9324 | AN0573 | AN1215 |
| | | AN8939 | AN6132 | AN8428 |
| | | AN2343 | AN6633 | AN3017 |
| | | AN9347 | AN10821 | AN2509 |
| | | AN7229 | AN10951 | AN7128 |
| | | AN3218 | AN6833 | AN6381 |
| | | AN3175 | AN0196 | AN1279 |
| | | AN1781 | AN8383 | AN1677 |
| | | AN3762 | AN2614 | AN8459 |
| | | AN2833 | AN7989 | AN7693 |
| | | AN5781 | AN1112 | AN2041 |
| | | AN7709 | AN1445 | AN9476 |
| | | AN2177 | AN2060 | AN3801 |
| | | AN7116 | AN8326 | AN2959 |
| | | AN3229 | AN11478 | AN3204 |
| | | AN3612 | AN6791 | AN8328 |
| | | AN5936 | AN4525 | AN0365 |
| | | AN2607 | AN8492 | AN2352 |
| | | AN1586 | AN3325 | AN6966 |
| | | AN1612 | AN8981 | AN2379 |
| | | AN4049 | AN11289 | AN8666 |
| | | AN2605 | AN3727 | AN2357 |
| | | AN6748 | AN8438 | AN6967 |
| | | AN0513 | AN0805 | AN5355 |
| | | AN9252 | AN9260 | AN8082 |
| | | AN2793 | AN0508 | AN3533 |
| | | AN9182 | AN11094 | AN1008 |
| | | AN7708 | AN6940 | AN8939 |
| | | AN4806 | AN0799 | AN9347 |
| | | AN7378 | AN8919 | AN4482 |
| | | AN0501 | AN2935 | AN2343 |
| | | AN6273 | AN9042 | AN7229 |
| | | AN5357 | AN3747 | AN7179 |
| | | AN1036 | AN9344 | AN5073 |
| | | AN0500 | AN1396 | AN0023 |

| | | | | |
|---|---|---|---|---|
| | | AN8471<br>AN2821<br>AN8121<br>AN8950<br>AN6963<br>AN7023<br>AN3130<br>AN7796<br>AN7955<br>AN7323<br>AN2960<br>AN5480<br>AN8303<br>AN0467<br>AN7753<br>AN2585<br>AN7809<br>AN8220<br>AN9432<br>AN7878<br>AN9202<br>AN2915<br>AN1816<br>AN8354<br>AN6386<br>AN3971<br>AN5459<br>AN1645<br>AN1320<br>AN0652<br>AN2806<br>AN2659<br>AN8608<br>AN5319<br>AN3560<br>AN3256<br>AN6103<br>AN3242<br>AN7703<br>AN2118<br>AN8090<br>AN3960<br>AN0510<br>AN5051<br>AN6401<br>AN2798<br>AN5017<br>AN6969<br>AN7401<br>AN9031<br>AN1703<br>AN5023<br>AN4147<br>AN2560<br>AN5823<br>AN4862<br>AN3562<br>AN8003<br>AN6414<br>AN8949<br>AN2610<br>AN9191<br>AN0768<br>AN4376<br>AN2827<br>AN0008 | AN11622<br>AN1585<br>AN8624<br>AN8441<br>AN4507<br>AN3334<br>AN6771<br>AN8195<br>AN6384<br>AN8275<br>AN6765<br>AN8138<br>AN2182<br>AN1813<br>AN7215<br>AN4198<br>AN0916<br>AN5304<br>AN0932<br>AN11050<br>AN11214<br>AN0099<br>AN11300<br>AN5545<br>AN10109<br>AN2619<br>AN11523<br>AN7404<br>AN0356<br>AN10422<br>AN2790<br>AN5424<br>AN8384<br>AN8905<br>AN5887<br>AN2037<br>AN7937<br>AN3769<br>AN4573<br>AN2695<br>AN2875<br>AN0743<br>AN2618<br>AN10291<br>AN1567<br>AN0858<br>AN0177<br>AN1499<br>AN7406<br>AN6776<br>AN9010<br>AN8803<br>AN0620<br>AN8205<br>AN8733<br>AN5891<br>AN10391<br>AN10108<br>AN4379<br>AN8134<br>AN5466<br>AN3246<br>AN6249<br>AN10913<br>AN5886<br>AN0927 | AN1781<br>AN2833<br>AN9299<br>AN3762<br>AN3043<br>AN7709<br>AN7265<br>AN7211<br>AN3674<br>AN8964<br>AN1612<br>AN5140<br>AN2793<br>AN9182<br>AN9166<br>AN1304<br>AN7708<br>AN4806<br>AN6761<br>AN8737<br>AN7051<br>AN0500<br>AN7269<br>AN7175<br>AN6581<br>AN8121<br>AN7023<br>AN6963<br>AN3130<br>AN7955<br>AN1032<br>AN8971<br>AN2960<br>AN0993<br>AN8619<br>AN9202<br>AN8597<br>AN7809<br>AN2915<br>AN9432<br>AN1816<br>AN7753<br>AN5465<br>AN3971<br>AN5459<br>AN3610<br>AN9454<br>AN3560<br>AN2806<br>AN7136<br>AN2659<br>AN6103<br>AN3805<br>AN5085<br>AN7058<br>AN7703<br>AN6011<br>AN8090<br>AN2118<br>AN0457<br>AN7790<br>AN9183<br>AN3415<br>AN5650<br>AN7129<br>AN7130 |

| | | | | |
|---|---|---|---|---|
| | | AN8764<br>AN2832<br>AN8113<br>AN7983<br>AN8400<br>AN7622<br>AN6408<br>AN1651<br>AN0100<br>AN2120<br>AN5037<br>AN7112<br>AN7024<br>AN0170<br>AN4363<br>AN3523<br>AN8996<br>AN4583<br>AN3052<br>AN8473<br>AN9280<br>AN5652<br>AN7204<br>AN1592<br>AN3400<br>AN0492<br>AN7825<br>AN9337<br>AN4367<br>AN2901<br>AN2488<br>AN7780<br>AN0036<br>AN8309<br>AN6657<br>AN8558<br>AN0915<br>AN7827<br>AN7959<br>AN4669<br>AN1214<br>AN2542<br>AN8570<br>AN5371<br>AN5893<br>AN3889<br>AN2530<br>AN4148<br>AN8199<br>AN9012<br>AN2564<br>AN9054<br>AN2846<br>AN3304<br>AN8312<br>AN6658<br>AN9343<br>AN8989<br>AN4590<br>AN8530<br>AN3327<br>AN5092<br>AN4029<br>AN8595<br>AN8425<br>AN7621 | AN9493<br>AN4199<br>AN11073<br>AN7529<br>AN5379<br>AN9186<br>AN1338<br>AN10015<br>AN1020<br>AN2840<br>AN3507<br>AN7851<br>AN11321<br>AN2839<br>AN4604<br>AN0197<br>AN10499<br>AN1726<br>AN3548<br>AN4545<br>AN11434<br>AN4913<br>AN7240<br>AN8891<br>AN11379<br>AN6781<br>AN3235<br>AN11029<br>AN5615<br>AN8357<br>AN7954<br>AN9508<br>AN3337<br>AN8992<br>AN11176<br>AN2587<br>AN1313<br>AN2830<br>AN1532<br>AN3298<br>AN8752<br>AN9196<br>AN10447<br>AN5273<br>AN7235<br>AN0588<br>AN7999<br>AN11459<br>AN9148<br>AN2321<br>AN7862<br>AN11479<br>AN11278<br>AN1246<br>AN8985<br>AN3480<br>AN8729<br>AN6274<br>AN7904<br>AN7087<br>AN1858<br>AN7365<br>AN1552<br>AN6746<br>AN4801<br>AN9377 | AN1703<br>AN6969<br>AN3773<br>AN5023<br>AN4147<br>AN8892<br>AN7523<br>AN2560<br>AN5823<br>AN4862<br>AN9279<br>AN8003<br>AN6414<br>AN0768<br>AN8740<br>AN4376<br>AN0008<br>AN8764<br>AN9478<br>AN8482 |

| | | | |
|---|---|---|---|
| | | AN4051 | AN9005 |
| | | AN5636 | AN4094 |
| | | AN8314 | AN5487 |
| | | AN2951 | AN9538 |
| | | AN2813 | AN7683 |
| | | AN2196 | AN7865 |
| | | AN2924 | AN11398 |
| | | AN7226 | AN2398 |
| | | AN7828 | AN8689 |
| | | AN2750 | AN3008 |
| | | AN8398 | AN0534 |
| | | AN8639 | AN9285 |
| | | AN8933 | AN7099 |
| | | AN3884 | AN6075 |
| | | AN8299 | AN7154 |
| | | AN1616 | AN10035 |
| | | AN4132 | AN9311 |
| | | AN8019 | AN7230 |
| | | AN4829 | AN0709 |
| | | AN2384 | AN3270 |
| | | AN9189 | AN6058 |
| | | AN3257 | AN5361 |
| | | AN0035 | AN2287 |
| | | AN7383 | AN5207 |
| | | AN5261 | AN6792 |
| | | AN8365 | AN11172 |
| | | AN0749 | AN11137 |
| | | AN7875 | AN10325 |
| | | AN2603 | AN1153 |
| | | AN1787 | AN10736 |
| | | AN8453 | AN2893 |
| | | AN8592 | AN6778 |
| | | AN8149 | AN10630 |
| | | AN8221 | AN8444 |
| | | AN7079 | AN0669 |
| | | AN0397 | AN7779 |
| | | AN3132 | AN6292 |
| | | AN2191 | AN3980 |
| | | AN9177 | AN4792 |
| | | AN0726 | AN2043 |
| | | AN8342 | AN2954 |
| | | AN8446 | AN2580 |
| | | AN7962 | AN9439 |
| | | AN2864 | AN8306 |
| | | AN8540 | AN7524 |
| | | AN7832 | AN5071 |
| | | AN9165 | AN0472 |
| | | AN5393 | AN3030 |
| | | AN7932 | AN6448 |
| | | AN4330 | AN9244 |
| | | AN8545 | AN4145 |
| | | AN3272 | AN6717 |
| | | AN0528 | AN4681 |
| | | AN9129 | AN0394 |
| | | AN3570 | AN7241 |
| | | AN3233 | AN3300 |
| | | AN6881 | AN10634 |
| | | AN0393 | AN5994 |
| | | AN4129 | AN3403 |
| | | AN1378 | AN0045 |
| | | AN5437 | AN2375 |
| | | AN3244 | AN8544 |
| | | AN8648 | AN3521 |
| | | AN8999 | AN0485 |
| | | AN9055 | AN1596 |
| | | AN4975 | AN7280 |

| | | | |
|---|---|---|---|
| | | AN7939 | AN7805 |
| | | AN3573 | AN0362 |
| | | AN9296 | AN7947 |
| | | AN1615 | AN2030 |
| | | AN6659 | AN3424 |
| | | AN2576 | AN2369 |
| | | AN8115 | AN2831 |
| | | AN8685 | AN3627 |
| | | AN3207 | AN7414 |
| | | AN7061 | AN7530 |
| | | AN1144 | AN8982 |
| | | AN3079 | AN6431 |
| | | AN3078 | AN9178 |
| | | AN8640 | AN8235 |
| | | AN9197 | AN7517 |
| | | AN2336 | AN2504 |
| | | AN2201 | AN11024 |
| | | AN3109 | AN7902 |
| | | AN8773 | AN11612 |
| | | AN7181 | AN11048 |
| | | AN0495 | AN10051 |
| | | AN2571 | AN11341 |
| | | AN8130 | AN6134 |
| | | AN1185 | AN7170 |
| | | AN9046 | AN9379 |
| | | AN0403 | AN0330 |
| | | AN5218 | AN7973 |
| | | AN8563 | AN9353 |
| | | AN9262 | AN3225 |
| | | AN2562 | AN2465 |
| | | AN9123 | AN3667 |
| | | AN1620 | AN8568 |
| | | AN9309 | AN5426 |
| | | AN1543 | AN7394 |
| | | AN3314 | AN7167 |
| | | AN8974 | AN7042 |
| | | AN7151 | AN7212 |
| | | AN7880 | AN7078 |
| | | AN4815 | AN7059 |
| | | AN6833 | AN3707 |
| | | AN8105 | AN10819 |
| | | AN0748 | AN10767 |
| | | AN5378 | AN1549 |
| | | AN7989 | AN4809 |
| | | AN0527 | AN4110 |
| | | AN2468 | AN4946 |
| | | AN2600 | AN9154 |
| | | AN4999 | AN7033 |
| | | AN8326 | AN5421 |
| | | AN8327 | AN8316 |
| | | AN3325 | AN9364 |
| | | AN8981 | AN1828 |
| | | AN7251 | AN2598 |
| | | AN8438 | AN7801 |
| | | AN0508 | AN3388 |
| | | AN1309 | AN11012 |
| | | AN7884 | AN6028 |
| | | AN6664 | AN8041 |
| | | AN0799 | AN11079 |
| | | AN8155 | AN0795 |
| | | AN4106 | AN0477 |
| | | AN6057 | AN10818 |
| | | AN2319 | AN4190 |
| | | AN9341 | AN3537 |
| | | AN9042 | AN6417 |
| | | AN8651 | AN6803 |

| | | | |
|---|---|---|---|
| | | AN3557 | AN9350 |
| | | AN5477 | AN4849 |
| | | AN9219 | AN8053 |
| | | AN0644 | AN2362 |
| | | AN7488 | AN3211 |
| | | AN9013 | AN6358 |
| | | AN1585 | AN3215 |
| | | AN8624 | AN8200 |
| | | AN3163 | AN10320 |
| | | AN4507 | AN2371 |
| | | AN0618 | AN2537 |
| | | AN5041 | AN10176 |
| | | AN6315 | AN2010 |
| | | AN6153 | AN2871 |
| | | AN3334 | AN7928 |
| | | AN6384 | AN5726 |
| | | AN8479 | AN8623 |
| | | AN6765 | AN1618 |
| | | AN8451 | AN2575 |
| | | AN0231 | AN3414 |
| | | AN7215 | AN3977 |
| | | AN8008 | AN11288 |
| | | AN1813 | AN4103 |
| | | AN8150 | AN11391 |
| | | AN0343 | AN2410 |
| | | AN3987 | AN10389 |
| | | AN7972 | AN0606 |
| | | AN2698 | AN0913 |
| | | AN6535 | AN5256 |
| | | AN4888 | AN3041 |
| | | AN3472 | AN5292 |
| | | AN5304 | AN6397 |
| | | AN9277 | AN8370 |
| | | AN8460 | AN11350 |
| | | AN4924 | AN3219 |
| | | AN9345 | AN2161 |
| | | AN3243 | AN7148 |
| | | AN2826 | AN11511 |
| | | AN7592 | AN7892 |
| | | AN5545 | AN6666 |
| | | AN2325 | AN7897 |
| | | AN5976 | AN8442 |
| | | AN0899 | AN9171 |
| | | AN6576 | AN6618 |
| | | AN5841 | AN11447 |
| | | AN5424 | AN10655 |
| | | AN6789 | AN1740 |
| | | AN2228 | AN6481 |
| | | AN8376 | AN3981 |
| | | AN8905 | AN7649 |
| | | AN3248 | AN8739 |
| | | AN3214 | AN7500 |
| | | AN0400 | AN8472 |
| | | AN2037 | AN1308 |
| | | AN9457 | AN8419 |
| | | AN7237 | AN5906 |
| | | AN0364 | AN10895 |
| | | AN3769 | AN2009 |
| | | AN3915 | AN3804 |
| | | AN8141 | AN7368 |
| | | AN1738 | AN0422 |
| | | AN2695 | AN4324 |
| | | AN2875 | AN7906 |
| | | AN9314 | AN6832 |
| | | AN8269 | AN10901 |
| | | AN0973 | AN2033 |

| | | | |
|---|---|---|---|
| | | AN3200<br>AN6648<br>AN5956<br>AN3551<br>AN2618<br>AN8422<br>AN1567<br>AN0548<br>AN7908<br>AN0858<br>AN7406<br>AN0169<br>AN8439<br>AN1786<br>AN8803<br>AN0332<br>AN3967<br>AN8134<br>AN7911<br>AN5466<br>AN6398<br>AN2200<br>AN6249<br>AN0418<br>AN2591<br>AN6399<br>AN5272<br>AN8612<br>AN8611<br>AN4769<br>AN6766<br>AN5090<br>AN4257<br>AN7022<br>AN5651<br>AN0015<br>AN7851<br>AN3507<br>AN5354<br>AN8262<br>AN7949<br>AN4604<br>AN5935<br>AN3548<br>AN4485<br>AN6847<br>AN6781<br>AN5427<br>AN3235<br>AN3504<br>AN2404<br>AN8093<br>AN5286<br>AN5615<br>AN3260<br>AN7954<br>AN1952<br>AN8952<br>AN2192<br>AN7408<br>AN3951<br>AN8992<br>AN2587<br>AN5089<br>AN2583<br>AN8131 | AN1419<br>AN8413<br>AN3717<br>AN6297<br>AN7788<br>AN11267<br>AN8319<br>AN11122<br>AN1148<br>AN8889<br>AN10779<br>AN10657<br>AN10097<br>AN1842<br>AN1605<br>AN6485<br>AN0198<br>AN5159<br>AN2697<br>AN4355<br>AN11206<br>AN10930<br>AN11196<br>AN10580<br>AN11451<br>AN0136<br>AN6298<br>AN7616<br>AN5945<br>AN3329<br>AN10083<br>AN10811<br>AN2661<br>AN5305<br>AN8921<br>AN4845<br>AN11201<br>AN2175<br>AN2561<br>AN11316<br>AN1647<br>AN2710<br>AN0484<br>AN5471<br>AN6697<br>AN11310<br>AN8362<br>AN11217<br>AN10506<br>AN8928<br>AN0693<br>AN3315<br>AN7874<br>AN6456<br>AN1930<br>AN2064<br>AN11149<br>AN7716<br>AN10409<br>AN6005<br>AN3866<br>AN4963<br>AN6918<br>AN7243<br>AN3563<br>AN5031 |

| | | | |
|---|---|---|---|
| | | AN4770 | AN1767 |
| | | AN9490 | AN1646 |
| | | AN0941 | AN2472 |
| | | AN7095 | AN3725 |
| | | AN3298 | AN11580 |
| | | AN1532 | AN6040 |
| | | AN3285 | AN10334 |
| | | AN2572 | AN1329 |
| | | AN1314 | AN11011 |
| | | AN7088 | AN1113 |
| | | AN5273 | AN3957 |
| | | AN5029 | AN5295 |
| | | AN1312 | AN11595 |
| | | AN8538 | AN7636 |
| | | AN0588 | AN10659 |
| | | AN3558 | AN10914 |
| | | AN3344 | AN2717 |
| | | AN2412 | AN10920 |
| | | AN7999 | AN5559 |
| | | AN3514 | AN4964 |
| | | AN3384 | AN6409 |
| | | AN1887 | AN4826 |
| | | AN2374 | AN6016 |
| | | AN2321 | AN8813 |
| | | AN9056 | AN8095 |
| | | AN1318 | AN0474 |
| | | AN8941 | AN11658 |
| | | AN9213 | AN7225 |
| | | AN0488 | AN1301 |
| | | AN5616 | AN10293 |
| | | AN8985 | AN0029 |
| | | AN3321 | AN2648 |
| | | AN7836 | AN2436 |
| | | AN8449 | AN7737 |
| | | AN5964 | AN3714 |
| | | AN2869 | AN6719 |
| | | AN5929 | AN2365 |
| | | AN8642 | AN8178 |
| | | AN4436 | AN11183 |
| | | AN1858 | AN7750 |
| | | AN9243 | AN10646 |
| | | AN5975 | AN3880 |
| | | AN4909 | AN3829 |
| | | AN8136 | AN6320 |
| | | AN7841 | AN0005 |
| | | AN3238 | AN3254 |
| | | AN0456 | AN1804 |
| | | AN1317 | AN11468 |
| | | AN8302 | AN0734 |
| | | AN9005 | AN0662 |
| | | AN5487 | AN5433 |
| | | AN4094 | AN9500 |
| | | AN6509 | AN1595 |
| | | AN0660 | AN10600 |
| | | AN5019 | AN0012 |
| | | AN7075 | AN7952 |
| | | AN7683 | AN8431 |
| | | AN4628 | AN6796 |
| | | AN9301 | AN8636 |
| | | AN7865 | AN0149 |
| | | AN0208 | AN0895 |
| | | AN2398 | AN6921 |
| | | AN8689 | AN8554 |
| | | AN3008 | AN6151 |
| | | AN7549 | AN11189 |
| | | AN0534 | AN8189 |

| | | | |
|---|---|---|---|
| | | AN9285 | AN7942 |
| | | AN8341 | AN1918 |
| | | AN1692 | AN6965 |
| | | AN7263 | AN7917 |
| | | AN8657 | AN2720 |
| | | AN2040 | AN6019 |
| | | AN7900 | AN1815 |
| | | AN7154 | AN2392 |
| | | AN9311 | AN2932 |
| | | AN1556 | AN6422 |
| | | AN0709 | AN1493 |
| | | AN6944 | AN0773 |
| | | AN6058 | AN7863 |
| | | AN5361 | AN0341 |
| | | AN8408 | AN1827 |
| | | AN1631 | AN1269 |
| | | AN6792 | AN10969 |
| | | AN5226 | AN7232 |
| | | AN1899 | AN9209 |
| | | AN7063 | AN10113 |
| | | AN7018 | AN6484 |
| | | AN7532 | AN7692 |
| | | AN8632 | AN3239 |
| | | AN0014 | AN11590 |
| | | AN7818 | AN0006 |
| | | AN1153 | AN3532 |
| | | AN2800 | AN8169 |
| | | AN0223 | AN4119 |
| | | AN5885 | AN7092 |
| | | AN8444 | AN7957 |
| | | AN7392 | AN2921 |
| | | AN3223 | AN5472 |
| | | AN8348 | AN1414 |
| | | AN6292 | AN5308 |
| | | AN3980 | AN5410 |
| | | AN7264 | AN6084 |
| | | AN2954 | AN7247 |
| | | AN9439 | AN9267 |
| | | AN0798 | AN4539 |
| | | AN5568 | AN1553 |
| | | AN4173 | AN3538 |
| | | AN5071 | AN6996 |
| | | AN9128 | AN10237 |
| | | AN8692 | AN2377 |
| | | AN2656 | AN4577 |
| | | AN0694 | AN8007 |
| | | AN1928 | AN9158 |
| | | AN5213 | AN8157 |
| | | AN9244 | AN4336 |
| | | AN7518 | AN7883 |
| | | AN2799 | AN3863 |
| | | AN7854 | AN4527 |
| | | AN0546 | AN11499 |
| | | AN7241 | AN6393 |
| | | AN3333 | AN6871 |
| | | AN7882 | AN10034 |
| | | AN3553 | AN5083 |
| | | AN5368 | AN11276 |
| | | AN5665 | AN3306 |
| | | AN5289 | AN10507 |
| | | AN3403 | AN6729 |
| | | AN9094 | AN6870 |
| | | AN2796 | AN6917 |
| | | AN5270 | AN4952 |
| | | AN8347 | AN3485 |
| | | AN7970 | AN10836 |

| | | | |
|---|---|---|---|
| | | AN2127 | AN10299 |
| | | AN3139 | AN2376 |
| | | AN2375 | AN10426 |
| | | AN2403 | AN11504 |
| | | AN8544 | AN10603 |
| | | AN6942 | AN7861 |
| | | AN3521 | AN5309 |
| | | AN2624 | AN6324 |
| | | AN7054 | AN11209 |
| | | AN7452 | AN10224 |
| | | AN6753 | AN10041 |
| | | AN9297 | AN10877 |
| | | AN7805 | AN2116 |
| | | AN5015 | AN6973 |
| | | AN2030 | AN3053 |
| | | AN3289 | AN5069 |
| | | AN7823 | AN10419 |
| | | AN8943 | AN8114 |
| | | AN7415 | AN8664 |
| | | AN8582 | AN8958 |
| | | AN2369 | AN5447 |
| | | AN0737 | AN5454 |
| | | AN8778 | AN7388 |
| | | AN5093 | AN9032 |
| | | AN6359 | AN7008 |
| | | AN2831 | AN11615 |
| | | AN3627 | AN7953 |
| | | AN7414 | AN3433 |
| | | AN3858 | AN8369 |
| | | AN8982 | AN5430 |
| | | AN5276 | AN8522 |
| | | AN6167 | AN4006 |
| | | AN6089 | AN10085 |
| | | AN7635 | AN0511 |
| | | AN3255 | AN10932 |
| | | AN8552 | AN9393 |
| | | AN0531 | AN11510 |
| | | AN9178 | AN6003 |
| | | AN7176 | AN1261 |
| | | AN5370 | AN8986 |
| | | AN1623 | AN7073 |
| | | AN7517 | AN2894 |
| | | AN1737 | AN9256 |
| | | AN6356 | AN8466 |
| | | AN3476 | AN6958 |
| | | AN1574 | AN8714 |
| | | AN9303 | AN9336 |
| | | AN5558 | AN8769 |
| | | AN7070 | AN4164 |
| | | AN7867 | AN4970 |
| | | AN0016 | AN5581 |
| | | AN9351 | AN8477 |
| | | AN6936 | AN1637 |
| | | AN7170 | AN1241 |
| | | AN6751 | AN9313 |
| | | AN6134 | AN6169 |
| | | AN1664 | AN2099 |
| | | AN7391 | AN1188 |
| | | AN0330 | AN0549 |
| | | AN7973 | AN5864 |
| | | AN9353 | AN4155 |
| | | AN2558 | AN2346 |
| | | AN3361 | AN3870 |
| | | AN1322 | AN1853 |
| | | AN7910 | AN4640 |
| | | AN5426 | AN2843 |

| | | | |
|---|---|---|---|
| | | AN2950<br>AN7394<br>AN7167<br>AN0056<br>AN7212<br>AN5507<br>AN1262<br>AN7813<br>AN2679<br>AN9332<br>AN7059<br>AN4854<br>AN8324<br>AN1549<br>AN6671<br>AN6959<br>AN4110<br>AN7842<br>AN0958<br>AN3531<br>AN2550<br>AN5053<br>AN3336<br>AN9044<br>AN3985<br>AN9364<br>AN8316<br>AN5244<br>AN8955<br>AN7801<br>AN2598<br>AN2198<br>AN3396<br>AN2340<br>AN8160<br>AN3574<br>AN5161<br>AN3150<br>AN3388<br>AN1882<br>AN0395<br>AN5435<br>AN5068<br>AN0020<br>AN4627<br>AN5938<br>AN2675<br>AN9350<br>AN3420<br>AN5417<br>AN2320<br>AN6358<br>AN0946<br>AN9317<br>AN8514<br>AN8962<br>AN4711<br>AN2010<br>AN9220<br>AN3414<br>AN1741<br>AN5937<br>AN8983<br>AN1582<br>AN7961<br>AN7281 | AN3127<br>AN3786<br>AN11245<br>AN10313<br>AN1302<br>AN2507<br>AN1007<br>AN6741<br>AN2249<br>AN11223<br>AN6385<br>AN1145<br>AN7142<br>AN7877<br>AN9302<br>AN11356<br>AN7756<br>AN2445<br>AN7767<br>AN7967<br>AN9503<br>AN2024<br>AN2031<br>AN11274<br>AN7800<br>AN11602<br>AN8358<br>AN2810<br>AN6255<br>AN3961<br>AN3820<br>AN5021<br>AN1460<br>AN1278<br>AN3059<br>AN10488<br>AN8127<br>AN5762<br>AN9163<br>AN10285<br>AN10533<br>AN7153<br>AN10326<br>AN4208<br>AN5284<br>AN11170<br>AN10251<br>AN10845<br>AN11368<br>AN0337<br>AN1749<br>AN2467<br>AN4852<br>AN10380<br>AN6723<br>AN1030<br>AN3195<br>AN7409<br>AN8591<br>AN7792<br>AN4125<br>AN10752<br>AN2540<br>AN7531<br>AN8166<br>AN6346 |

| | | | |
|---|---|---|---|
| | | AN1644<br>AN5544<br>AN8970<br>AN5831<br>AN2004<br>AN0102<br>AN3041<br>AN6397<br>AN9206<br>AN5444<br>AN8370<br>AN8617<br>AN6121<br>AN7082<br>AN5110<br>AN8630<br>AN8470<br>AN3406<br>AN0158<br>AN7171<br>AN9171<br>AN7897<br>AN8442<br>AN3713<br>AN7879<br>AN8364<br>AN2039<br>AN8779<br>AN1630<br>AN6873<br>AN8739<br>AN6718<br>AN9152<br>AN1835<br>AN2254<br>AN9205<br>AN7278<br>AN5664<br>AN3273<br>AN0509<br>AN0750<br>AN4807<br>AN7916<br>AN9348<br>AN7368<br>AN4195<br>AN0601<br>AN7267<br>AN0024<br>AN7906<br>AN9139<br>AN7647<br>AN9203<br>AN2033<br>AN1450<br>AN1742<br>AN1591<br>AN7152<br>AN7788<br>AN4642<br>AN1903<br>AN2814<br>AN1605<br>AN3475<br>AN6947<br>AN4621 | AN11156<br>AN3879<br>AN6801<br>AN1826<br>AN6362<br>AN3881<br>AN9535<br>AN8366<br>AN3813<br>AN5473<br>AN1206<br>AN6941<br>AN3522<br>AN8620<br>AN0265<br>AN2712<br>AN2034<br>AN4065<br>AN1193<br>AN9259<br>AN9331<br>AN0007<br>AN1832<br>AN5930<br>AN9338<br>AN7396<br>AN7866<br>AN4373<br>AN7279<br>AN7120<br>AN8106<br>AN11645<br>AN5546<br>AN3761<br>AN7053<br>AN10025<br>AN7590<br>AN3410<br>AN1580<br>AN5973<br>AN7157<br>AN11269<br>AN1914<br>AN5564<br>AN8241<br>AN3994<br>AN10931<br>AN7387<br>AN0742<br>AN10505<br>AN2804<br>AN5135<br>AN9315<br>AN3102<br>AN5422<br>AN11569<br>AN5902<br>AN3284<br>AN6804<br>AN9168<br>AN6104<br>AN10136<br>AN8416<br>AN10364<br>AN5065<br>AN0640 |

| | | | |
|---|---|---|---|
| | | AN3252<br>AN2697<br>AN2928<br>AN2411<br>AN5844<br>AN9427<br>AN6234<br>AN8433<br>AN4128<br>AN1203<br>AN6809<br>AN5334<br>AN9121<br>AN5945<br>AN3329<br>AN7888<br>AN4357<br>AN8338<br>AN4146<br>AN3986<br>AN3496<br>AN8921<br>AN1895<br>AN0214<br>AN6693<br>AN2710<br>AN2175<br>AN2561<br>AN1647<br>AN0515<br>AN2330<br>AN2199<br>AN9456<br>AN7089<br>AN2310<br>AN8587<br>AN3494<br>AN8494<br>AN2197<br>AN3253<br>AN7896<br>AN4506<br>AN7577<br>AN3315<br>AN0693<br>AN7874<br>AN2795<br>AN3282<br>AN7011<br>AN2064<br>AN1098<br>AN3479<br>AN7191<br>AN5285<br>AN3316<br>AN6005<br>AN9266<br>AN0283<br>AN5509<br>AN8610<br>AN3563<br>AN7398<br>AN2694<br>AN8108<br>AN2595<br>AN3023 | AN1837<br>AN9378<br>AN7322<br>AN4480<br>AN2687<br>AN2344<br>AN10303<br>AN11075<br>AN2664<br>AN4394<br>AN9026<br>AN3077<br>AN11031<br>AN5418<br>AN3159<br>AN2498<br>AN11161<br>AN3287<br>AN7850<br>AN1925<br>AN4870<br>AN5081<br>AN1784<br>AN6784<br>AN11628<br>AN6208<br>AN8353<br>AN5359<br>AN8381<br>AN1817<br>AN6056<br>AN1542<br>AN5842<br>AN8537<br>AN2599<br>AN2315<br>AN5310<br>AN2277<br>AN3250<br>AN5859<br>AN2225<br>AN5582<br>AN7203<br>AN8349<br>AN8333<br>AN0908<br>AN3027<br>AN8011<br>AN8234<br>AN8135<br>AN10581<br>AN11324<br>AN0299<br>AN4089<br>AN8755<br>AN1803<br>AN4116<br>AN2544<br>AN6491<br>AN2493<br>AN7981<br>AN5467<br>AN2955<br>AN3543<br>AN8415<br>AN10933 |

| | | | |
|---|---|---|---|
| | | AN6487<br>AN8927<br>AN2722<br>AN1614<br>AN9200<br>AN0600<br>AN5072<br>AN0959<br>AN1646<br>AN3545<br>AN1594<br>AN2730<br>AN1540<br>AN1307<br>AN1608<br>AN9242<br>AN8158<br>AN0974<br>AN7600<br>AN8104<br>AN7268<br>AN4474<br>AN7702<br>AN3950<br>AN6929<br>AN6411<br>AN9001<br>AN9004<br>AN1941<br>AN9375<br>AN5559<br>AN6662<br>AN8143<br>AN6409<br>AN4826<br>AN0622<br>AN4460<br>AN2892<br>AN8095<br>AN5396<br>AN1938<br>AN2649<br>AN5555<br>AN7225<br>AN8109<br>AN0029<br>AN0635<br>AN8596<br>AN2648<br>AN3387<br>AN8947<br>AN7808<br>AN6739<br>AN5076<br>AN9304<br>AN3714<br>AN7066<br>AN6719<br>AN0342<br>AN2365<br>AN4117<br>AN2714<br>AN1073<br>AN6403<br>AN2590<br>AN7814 | AN6787<br>AN11469<br>AN6091<br>AN10144<br>AN3210<br>AN4325<br>AN7402<br>AN2622<br>AN10868<br>AN8725<br>AN1432<br>AN6518<br>AN0396<br>AN3048<br>AN3555<br>AN6377<br>AN8083<br>AN11175<br>AN7169<br>AN2611<br>AN2828<br>AN8352<br>AN2097<br>AN2683<br>AN1659<br>AN8987<br>AN10345<br>AN1332<br>AN8621<br>AN6455<br>AN5942<br>AN9342<br>AN11466<br>AN11006<br>AN6955<br>AN2823<br>AN1321<br>AN5549<br>AN4216<br>AN8655<br>AN2348<br>AN2669<br>AN1089<br>AN2366<br>AN8483<br>AN3051<br>AN8399<br>AN6043<br>AN5079<br>AN5144<br>AN8424<br>AN2632<br>AN11096<br>AN3151<br>AN2347<br>AN8375<br>AN10424<br>AN1462<br>AN5372<br>AN11105<br>AN5577<br>AN10094<br>AN8593<br>AN9122<br>AN9027<br>AN7876 |

| | | | |
|---|---|---|---|
| | | AN3271<br>AN8935<br>AN4879<br>AN7057<br>AN0005<br>AN2402<br>AN1804<br>AN5034<br>AN7080<br>AN8428<br>AN3017<br>AN7128<br>AN0012<br>AN8142<br>AN7952<br>AN6381<br>AN1279<br>AN6796<br>AN1607<br>AN8459<br>AN7693<br>AN2041<br>AN3801<br>AN2959<br>AN3204<br>AN8328<br>AN7585<br>AN0895<br>AN8554<br>AN7295<br>AN2379<br>AN7942<br>AN1918<br>AN4130<br>AN2720<br>AN8082<br>AN6019<br>AN2392<br>AN4482<br>AN7179<br>AN5073<br>AN7863<br>AN0023<br>AN9299<br>AN0535<br>AN0341<br>AN3043<br>AN1269<br>AN7265<br>AN9209<br>AN7232<br>AN7211<br>AN7692<br>AN3239<br>AN3674<br>AN0006<br>AN8964<br>AN4119<br>AN5140<br>AN2921<br>AN5410<br>AN1672<br>AN9166<br>AN9267<br>AN1304<br>AN1792 | AN1755<br>AN9291<br>AN6203<br>AN7798<br>AN5101<br>AN0032<br>AN10333<br>AN1833<br>AN11420<br>AN3791<br>AN3614<br>AN5812<br>AN9161<br>AN8410<br>AN9247<br>AN7412<br>AN11477<br>AN10039<br>AN7341<br>AN11387<br>AN7511<br>AN8532<br>AN0619<br>AN7071<br>AN8465<br>AN11261<br>AN4658<br>AN8006<br>AN8652<br>AN9318<br>AN7385<br>AN11184<br>AN2792<br>AN5056<br>AN0871<br>AN6177<br>AN8101<br>AN9075<br>AN5002<br>AN8476<br>AN5845<br>AN11218<br>AN10297<br>AN11624<br>AN3391<br>AN5315<br>AN11339<br>AN9321<br>AN1176<br>AN7938<br>AN6810<br>AN11088<br>AN0220<br>AN4140<br>AN8295<br>AN8641<br>AN11337<br>AN3473<br>AN7115<br>AN5550<br>AN7317<br>AN3998<br>AN10831<br>AN8102<br>AN5843<br>AN8646 |

| | | | |
|---|---|---|---|
| | | AN1808<br>AN6761<br>AN8737<br>AN7051<br>AN7870<br>AN7269<br>AN3538<br>AN8007<br>AN7175<br>AN6581<br>AN9158<br>AN8157<br>AN9172<br>AN4336<br>AN7883<br>AN8526<br>AN1032<br>AN8971<br>AN0993<br>AN8597<br>AN6871<br>AN5083<br>AN8619<br>AN6870<br>AN3286<br>AN0878<br>AN3485<br>AN3495<br>AN5465<br>AN3203<br>AN2376<br>AN3610<br>AN9454<br>AN7136<br>AN1617<br>AN3805<br>AN8052<br>AN5085<br>AN7058<br>AN6011<br>AN0217<br>AN0457<br>AN7790<br>AN9183<br>AN2116<br>AN3415<br>AN1593<br>AN5069<br>AN9276<br>AN5650<br>AN7129<br>AN3302<br>AN1077<br>AN8114<br>AN4079<br>AN7130<br>AN0552<br>AN3773<br>AN8958<br>AN3975<br>AN5239<br>AN8892<br>AN9032<br>AN7523<br>AN4481<br>AN9279 | AN11049<br>AN3086<br>AN0845<br>AN3775<br>AN6004<br>AN11554<br>AN10104<br>AN10870<br>AN1311<br>AN0230<br>AN10430<br>AN6420<br>AN10915<br>AN9002<br>AN1897<br>AN5604<br>AN7418<br>AN9043<br>AN0778<br>AN11220<br>AN5944<br>AN6852<br>AN0468<br>AN7238<br>AN1558<br>AN7956<br>AN3872<br>AN2588<br>AN8495<br>AN8351<br>AN6525<br>AN3753<br>AN6367<br>AN4039<br>AN10502<br>AN10250<br>AN8513<br>AN8638<br>AN6636<br>AN1841<br>AN8371<br>AN4206<br>AN6862<br>AN7139<br>AN6821<br>AN0499<br>AN7413<br>AN6745<br>AN1604<br>AN6117<br>AN3901<br>AN10238<br>AN3297<br>AN8601<br>AN10749<br>AN11265<br>AN8890<br>AN8717<br>AN10547<br>AN10481<br>AN8899<br>AN3742<br>AN6464<br>AN3022<br>AN2784<br>AN8756 |

| | | | |
|---|---|---|---|
| | | AN6788<br>AN7953<br>AN8740<br>AN3433<br>AN2779<br>AN7856<br>AN9478<br>AN8482 | AN9492<br>AN10500<br>AN4654<br>AN8132<br>AN11354<br>AN4696<br>AN7988<br>AN3641<br>AN9014<br>AN1186<br>AN11427<br>AN6935<br>AN0761<br>AN4144<br>AN3259<br>AN6582<br>AN6656<br>AN8389<br>AN11584<br>AN9180<br>AN9188<br>AN5316<br>AN2609<br>AN3952<br>AN5556<br>AN5584<br>AN6063<br>AN1057<br>AN4842<br>AN11119<br>AN2625<br>AN7384<br>AN8480<br>AN2276<br>AN4483<br>AN10160<br>AN6410<br>AN8165<br>AN0497<br>AN3732<br>AN0523<br>AN0521<br>AN11028<br>AN2568<br>AN4141<br>AN10080<br>AN11450<br>AN5075<br>AN8977<br>AN3382<br>AN5367<br>AN5790<br>AN8913<br>AN1078<br>AN2286<br>AN1571<br>AN10023<br>AN4647<br>AN7728<br>AN2545<br>AN5231<br>AN7421<br>AN8907<br>AN11227<br>AN11625<br>AN7081 |

| | | | |
|---|---|---|---|
| | | | AN6322<br>AN5377<br>AN8555<br>AN8910<br>AN2084<br>AN6092<br>AN0022<br>AN2471<br>AN8374<br>AN2696<br>AN0811<br>AN0420<br>AN9235<br>AN8018<br>AN9073<br>AN11298<br>AN0224<br>AN10846<br>AN5996<br>AN1621<br>AN7996<br>AN4112<br>AN1142<br>AN6624<br>AN4659<br>AN4152<br>AN3032<br>AN0541<br>AN9052<br>AN9036<br>AN4111<br>AN1852<br>AN3777<br>AN6744<br>AN4390<br>AN3781<br>AN7642<br>AN6475<br>AN5328<br>AN4022<br>AN3566<br>AN5324<br>AN4438<br>AN5611<br>AN11233<br>AN10667<br>AN7644<br>AN4120<br>AN6242<br>AN11393<br>AN8979<br>AN8622<br>AN0180<br>AN8447<br>AN6376<br>AN5838<br>AN7202<br>AN11530<br>AN7543<br>AN8091<br>AN4603<br>AN3492<br>AN7026<br>AN7098<br>AN1140<br>AN1386 |

| | | | |
|---|---|---|---|
| | | | AN5240<br>AN8458<br>AN11099<br>AN3891<br>AN4503<br>AN11294<br>AN6402<br>AN0050<br>AN7625<br>AN10976<br>AN8037<br>AN8397<br>AN9286<br>AN5763<br>AN6653<br>AN2811<br>AN3308<br>AN7390<br>AN0940<br>AN2601<br>AN8840<br>AN9324<br>AN5591<br>AN1825<br>AN10771<br>AN9480<br>AN11474<br>AN7218<br>AN5363<br>AN5781<br>AN2177<br>AN8978<br>AN3229<br>AN3045<br>AN5936<br>AN1586<br>AN2958<br>AN4354<br>AN6748<br>AN1250<br>AN9485<br>AN7378<br>AN0501<br>AN5764<br>AN5357<br>AN3700<br>AN8471<br>AN6770<br>AN9050<br>AN9323<br>AN5054<br>AN8950<br>AN10910<br>AN7186<br>AN7796<br>AN8303<br>AN7878<br>AN10168<br>AN8354<br>AN5513<br>AN3020<br>AN7950<br>AN5921<br>AN1320<br>AN3213<br>AN10688 |

| | | | |
|---|---|---|---|
| | | | AN8608<br>AN3256<br>AN3242<br>AN1648<br>AN10075<br>AN10958<br>AN6264<br>AN3960<br>AN8027<br>AN3381<br>AN7290<br>AN3311<br>AN3680<br>AN5051<br>AN5589<br>AN6401<br>AN2798<br>AN5017<br>AN10278<br>AN11008<br>AN9268<br>AN4911<br>AN1588<br>AN8184<br>AN11187<br>AN11668<br>AN8400<br>AN3397<br>AN6670<br>AN4897<br>AN8456<br>AN2693<br>AN5063<br>AN7024<br>AN0170<br>AN2844<br>AN3290<br>AN3161<br>AN4583<br>AN9187<br>AN8473<br>AN1592<br>AN10045<br>AN3400<br>AN2919<br>AN7864<br>AN0492<br>AN7825<br>AN6301<br>AN4367<br>AN3701<br>AN7509<br>AN2350<br>AN6596<br>AN2488<br>AN10337<br>AN11614<br>AN4153<br>AN11525<br>AN8309<br>AN11557<br>AN2527<br>AN3816<br>AN11297<br>AN0711<br>AN11039 |

| | | | |
|---|---|---|---|
| | | | AN6436<br>AN4669<br>AN5371<br>AN2291<br>AN10286<br>AN10369<br>AN4062<br>AN8111<br>AN9272<br>AN6658<br>AN9343<br>AN7641<br>AN9264<br>AN4590<br>AN5092<br>AN2711<br>AN5636<br>AN8314<br>AN4816<br>AN5282<br>AN2924<br>AN11078<br>AN9227<br>AN2750<br>AN9287<br>AN9037<br>AN4270<br>AN3125<br>AN4921<br>AN8299<br>AN11633<br>AN11608<br>AN10847<br>AN4829<br>AN8088<br>AN3257<br>AN0035<br>AN5261<br>AN2603<br>AN1787<br>AN8221<br>AN0397<br>AN2107<br>AN0949<br>AN9208<br>AN7832<br>AN10669<br>AN7477<br>AN3277<br>AN3348<br>AN8775<br>AN10556<br>AN5180<br>AN9129<br>AN11627<br>AN6908<br>AN4129<br>AN3244<br>AN1506<br>AN11601<br>AN3079<br>AN2336<br>AN0764<br>AN0518<br>AN2571<br>AN6029 |

| | | | |
|---|---|---|---|
| | | | AN9197<br>AN0976<br>AN8432<br>AN11464<br>AN2742<br>AN5565<br>AN8130<br>AN0399<br>AN7086<br>AN9123<br>AN1543<br>AN8974<br>AN6452<br>AN7151<br>AN1055<br>AN7880<br>AN9329<br>AN0748<br>AN10493<br>AN0527<br>AN2600<br>AN4999<br>AN8327<br>AN8414<br>AN1624<br>AN1087<br>AN8653<br>AN11581<br>AN7055<br>AN7884<br>AN6664<br>AN8990<br>AN8759<br>AN1001<br>AN11432<br>AN11246<br>AN3163<br>AN11585<br>AN0618<br>AN5041<br>AN5847<br>AN8815<br>AN11454<br>AN10935<br>AN8150<br>AN11286<br>AN0343<br>AN11159<br>AN1286<br>AN7138<br>AN8478<br>AN4888<br>AN7871<br>AN3472<br>AN5860<br>AN3995<br>AN3520<br>AN9345<br>AN11314<br>AN1160<br>AN3066<br>AN8043<br>AN11626<br>AN2684<br>AN11673<br>AN2228 |

| | | | |
|---|---|---|---|
| | | | AN6789<br>AN8122<br>AN8376<br>AN3493<br>AN8437<br>AN9457<br>AN10245<br>AN7237<br>AN3915<br>AN8141<br>AN1738<br>AN7246<br>AN11439<br>AN0368<br>AN8542<br>AN8269<br>AN0973<br>AN5956<br>AN6480<br>AN6648<br>AN3551<br>AN3690<br>AN10270<br>AN2912<br>AN7908<br>AN0526<br>AN8439<br>AN1786<br>AN9124<br>AN7911<br>AN6398<br>AN6735<br>AN8074<br>AN0418<br>AN8237<br>AN2591<br>AN6399<br>AN5272<br>AN5731<br>AN3632<br>AN5905<br>AN6760<br>AN3330<br>AN6766<br>AN5090<br>AN5358<br>AN2383<br>AN5460<br>AN8152<br>AN2103<br>AN10028<br>AN3504<br>AN9236<br>AN10529<br>AN2067<br>AN2404<br>AN3212<br>AN3260<br>AN5579<br>AN1952<br>AN11121<br>AN8952<br>AN2019<br>AN2847<br>AN5923<br>AN6846 |

| | | | |
|---|---|---|---|
| | | | AN8131<br>AN4770<br>AN9490<br>AN11348<br>AN3285<br>AN1314<br>AN7088<br>AN9034<br>AN5264<br>AN1312<br>AN2566<br>AN3558<br>AN6747<br>AN7117<br>AN8454<br>AN6883<br>AN2374<br>AN3514<br>AN8941<br>AN1982<br>AN9213<br>AN11195<br>AN9310<br>AN10458<br>AN4436<br>AN9243<br>AN11268<br>AN7654<br>AN11317<br>AN8136<br>AN0078<br>AN3160<br>AN1317<br>AN5563<br>AN5917<br>AN9257<br>AN0741<br>AN5374<br>AN4917<br>AN7700<br>AN8031<br>AN2631<br>AN4628<br>AN2373<br>AN6453<br>AN10572<br>AN6872<br>AN0314<br>AN11215<br>AN5217<br>AN7991<br>AN8657<br>AN11133<br>AN9225<br>AN0263<br>AN5883<br>AN10612<br>AN8462<br>AN3848<br>AN10776<br>AN1899<br>AN8663<br>AN7056<br>AN8417<br>AN5890<br>AN6731 |

| | | | |
|---|---|---|---|
| | | | AN2146<br>AN3404<br>AN3418<br>AN0223<br>AN3223<br>AN5885<br>AN3896<br>AN6915<br>AN8936<br>AN11285<br>AN4819<br>AN11363<br>AN5568<br>AN2656<br>AN3230<br>AN0694<br>AN2808<br>AN6945<br>AN3281<br>AN11259<br>AN11461<br>AN4722<br>AN2799<br>AN0106<br>AN0159<br>AN11095<br>AN3333<br>AN5368<br>AN10296<br>AN5289<br>AN7376<br>AN7101<br>AN10123<br>AN8163<br>AN5270<br>AN0747<br>AN7970<br>AN8347<br>AN7657<br>AN10860<br>AN6753<br>AN0408<br>AN9297<br>AN9164<br>AN5015<br>AN1814<br>AN9162<br>AN3540<br>AN8943<br>AN0207<br>AN7415<br>AN5585<br>AN3258<br>AN4990<br>AN6089<br>AN6167<br>AN11442<br>AN5668<br>AN6128<br>AN7765<br>AN6635<br>AN1737<br>AN6356<br>AN5291<br>AN5405<br>AN5558 |

| | | | |
|---|---|---|---|
| | | | AN9426<br>AN7929<br>AN2558<br>AN0584<br>AN0227<br>AN3361<br>AN4443<br>AN2791<br>AN4823<br>AN10487<br>AN2634<br>AN0056<br>AN10452<br>AN1262<br>AN4205<br>AN5995<br>AN1890<br>AN0991<br>AN8382<br>AN5194<br>AN8324<br>AN8894<br>AN3336<br>AN3985<br>AN5244<br>AN7521<br>AN7607<br>AN3574<br>AN5161<br>AN3150<br>AN7551<br>AN0395<br>AN5435<br>AN11550<br>AN11575<br>AN0020<br>AN10060<br>AN11378<br>AN5628<br>AN9354<br>AN0698<br>AN0451<br>AN0602<br>AN5927<br>AN9317<br>AN0454<br>AN1799<br>AN7403<br>AN0171<br>AN2673<br>AN3990<br>AN10576<br>AN2662<br>AN1741<br>AN8983<br>AN10886<br>AN8196<br>AN4041<br>AN5544<br>AN8970<br>AN11313<br>AN3307<br>AN2920<br>AN0102<br>AN6932<br>AN10078 |

| | | | |
|---|---|---|---|
| | | | AN9241<br>AN5444<br>AN11120<br>AN6121<br>AN6831<br>AN1492<br>AN8470<br>AN5461<br>AN7171<br>AN9041<br>AN2543<br>AN2156<br>AN8219<br>AN11556<br>AN8957<br>AN9367<br>AN8364<br>AN2342<br>AN6048<br>AN4192<br>AN3856<br>AN0839<br>AN11205<br>AN8577<br>AN9152<br>AN0021<br>AN2254<br>AN9205<br>AN7278<br>AN9348<br>AN3535<br>AN9203<br>AN3188<br>AN2702<br>AN4642<br>AN5004<br>AN6947<br>AN4621<br>AN2928<br>AN2411<br>AN5844<br>AN7072<br>AN1501<br>AN9251<br>AN6234<br>AN6928<br>AN2654<br>AN11086<br>AN5334<br>AN7936<br>AN7888<br>AN10745<br>AN1597<br>AN3986<br>AN7873<br>AN1895<br>AN3556<br>AN0214<br>AN10880<br>AN6693<br>AN4500<br>AN5874<br>AN6673<br>AN6407<br>AN0655<br>AN8587 |

| | | | |
|---|---|---|---|
| | | | AN5538<br>AN6241<br>AN8494<br>AN10305<br>AN4506<br>AN1163<br>AN7034<br>AN11071<br>AN7899<br>AN7062<br>AN1098<br>AN5285<br>AN3316<br>AN9266<br>AN0283<br>AN3885<br>AN10332<br>AN1917<br>AN1614<br>AN2595<br>AN2722<br>AN8009<br>AN0959<br>AN2730<br>AN10972<br>AN1307<br>AN1540<br>AN1608<br>AN1754<br>AN1886<br>AN0974<br>AN7324<br>AN8104<br>AN3950<br>AN6929<br>AN7656<br>AN11494<br>AN6411<br>AN2646<br>AN10748<br>AN6454<br>AN1108<br>AN9004<br>AN1941<br>AN2582<br>AN7596<br>AN1330<br>AN4356<br>AN11283<br>AN5173<br>AN2876<br>AN5970<br>AN9365<br>AN4940<br>AN5251<br>AN1579<br>AN8109<br>AN8947<br>AN7808<br>AN4515<br>AN7166<br>AN9304<br>AN0342<br>AN4423<br>AN0885<br>AN11253 |

| | | | |
|---|---|---|---|
| | | | AN3099<br>AN6403<br>AN8311<br>AN2590<br>AN11291<br>AN2402<br>AN1625<br>AN8816<br>AN7041<br>AN10135<br>AN5453<br>AN6812<br>AN8142<br>AN1607<br>AN0841<br>AN2709<br>AN7003<br>AN8168<br>AN7585<br>AN7295<br>AN10558<br>AN10100<br>AN4130<br>AN0529<br>AN11051<br>AN10678<br>AN11533<br>AN9169<br>AN10786<br>AN5275<br>AN7207<br>AN0535<br>AN2822<br>AN8581<br>AN4009<br>AN4118<br>AN4825<br>AN5255<br>AN7149<br>AN3359<br>AN11456<br>AN6538<br>AN3354<br>AN3982<br>AN2628<br>AN1672<br>AN10861<br>AN1792<br>AN1808<br>AN0979<br>AN10448<br>AN7870<br>AN5047<br>AN8991<br>AN9172<br>AN5649<br>AN6818<br>AN8526<br>AN10577<br>AN5928<br>AN7164<br>AN3490<br>AN3278<br>AN3286<br>AN0878<br>AN10428 |

| | | | |
|---|---|---|---|
| | | | AN3495<br>AN3115<br>AN3203<br>AN5468<br>AN7930<br>AN1617<br>AN8052<br>AN9192<br>AN3734<br>AN4010<br>AN0217<br>AN10974<br>AN6127<br>AN6306<br>AN5060<br>AN1593<br>AN9276<br>AN9306<br>AN0172<br>AN3302<br>AN10310<br>AN1077<br>AN11518<br>AN4079<br>AN2546<br>AN0552<br>AN6836<br>AN10021<br>AN11197<br>AN1200<br>AN3975<br>AN5239<br>AN5523<br>AN6445<br>AN4481<br>AN9137<br>AN6788<br>AN3383<br>AN2779<br>AN1670<br>AN7856 |

S3 – Lists of primers used in the qPCR study.

| | |
|---|---|
| BrlA-qPCR-Fwd | TCTCCTACACCAACTCCAACAA |
| BrlA-qPCR-Rvs | ATTCGTTCCTGCCCTTCC |
| AbaA-qPCR-Fwd | ACCCTCCAGTCCCAGTCC |
| AbaA-qPCR-Rvs | TCACCATCTTTCCCAGTATCC |
| WetA-qPCR-Fwd | GCTTCTCCCCTGGCTTGAT |
| WetA-qPCR-Rvs | TTCTGCTGCGTTCCTCATCT |
| histone2b-qPCR-Fwd | CACCCGGACACTGGTATCTC |
| histone2b-qPCR-Rvs | GAATACTTCGTAACGGCCTTGG |
| AN0010-qPCR-Fwd | CGGGCACTACAGGAAGGAG |
| AN0010-qPCR-Rvs | TATCGGCACAGGAAAACG |
| AN3247-qPCR-Fwd | GCTCTTCTACCCGCACCTC |
| AN3247-qPCR-Rvs | GATACACCACGCACAAACTCA |
| AN3304-qPCR-Fwd | GCTAAAATGCCCCGAGAC |
| AN3304-qPCR-Rvs | CCGTTCCCAAGACCAGAC |
| AN8595-qPCR-Fwd | GGAGGAGGAGAGGAGGGTAA |
| AN8595-qPCR-Rvs | CAGAGGAGTTGGATGGTGGT |
| AN8610-qPCR-Fwd | GCTTCAACAAGGAGACCTACG |
| AN8610-qPCR-Rvs | TGGACACGAACAGGACGAT |
| AN9165-qPCR-Fwd | CTCGTCATCTCCACCTACAGC |
| AN9165-qPCR-Rvs | GGACAAACTCGCACCACAG |
